# RV-specific Targeting of Snai1 Rescues Pulmonary Hypertension-induced Right Ventricular Failure by Inhibiting EndMT and Fibrosis *via* LOXL2 Mediated Mechanism

**DOI:** 10.1101/2024.04.30.591766

**Authors:** Somanshu Banerjee, Varina R. Clark Onwunyi, Jason Hong, Sandra Martineau, Gregory A. Fishbein, Sandra Breuils Bonnet, Steeve Provencher, Sébastien Bonnet, Soban Umar

## Abstract

**Background:** Pulmonary hypertension (PH)-induced right ventricular (RV) failure (PH-RVF) is a significant prognostic determinant of mortality and is characterized by RV hypertrophy, endothelial-to-mesenchymal transition (EndMT), fibroblast-to-myofibroblast transition (FMT), fibrosis, and extracellular matrix (ECM)-remodeling. Despite the importance of RV function in PH, the mechanistic details of PH-RVF, especially the regulatory control of RV EndMT, FMT, and fibrosis, remain unclear. The action of transcription factor Snai1 is shown to be mediated through LOXL2 recruitment, and their co-translocation to the nucleus, during EndMT progression. We hypothesize that RV EndMT and fibrosis in PH-RVF are governed by the TGFβ1-Snai1-LOXL2 axis. Furthermore, targeting Snai1 could serve as a novel therapeutic strategy for PH-RVF.

**Methods:** Adult male Sprague Dawley rats (250-300g) received either a single subcutaneous injection of Monocrotaline (MCT, 60mg/kg, n=9; followed for 30-days) or Sugen (SU5416 20mg/kg, n=9; 10% O_2_ hypoxia for 3-weeks followed by normoxia for 2-weeks) or PBS (CTRL, n=9). We performed secondary bioinformatics analysis on the RV bulk RNA-Seq data from MCT, SuHx, and PAB rats and human PH-PVF. We validated EndMT and FMT and their association with Snai1 and LOXL2 in the RVs of MCT and SuHx rat models and human PH-RVF using immunofluorescence, qPCR, and Western blots. For *in vivo* Snai1 knockdown (Snai1-KD), MCT-rats either received Snai1-siRNA (n=7; 5nM/injection every 3-4 days; 4-injections) or scramble (SCRM-KD; n=7) through tail vein from day 14-30 after MCT. Echocardiography and catheterization were performed terminally. Bulk RNASeq and differential expression analysis were performed on Snai1- and SCRM-KD rat RVs. *In vitro* Snai1-KD was performed on human coronary artery endothelial cells (HCAECs) and human cardiac fibroblasts (HCFs) under hypoxia+TGFβ1 for 72-hrs.

**Results:** PH-RVF had increased RVSP and Fulton index and decreased RV fractional area change (RVFAC %). RV RNASeq demonstrated EndMT as the common top-upregulated pathway between rat (MCT, SuHx, and PAB) and human PH-RVF. Immunofluorescence using EndMT- and FMT-specific markers demonstrated increased EndMT and FMT in RV of MCT and SuHx rats and PH-RVF patients. Further, RV expression of TGFβ1, Snai1, and LOXL2 was increased in MCT and SuHx. Nuclear co-localization and increased immunoreactivity, transcript, and protein levels of Snai1 and LOXL2 were observed in MCT and SuHx rats and human RVs. MCT rats treated with Snai1-siRNA demonstrated decreased Snai1 expression, RVSP, Fulton index, and increased RVFAC. Snai1-KD resulted in decreased RV-EndMT, FMT, and fibrosis *via* a LOXL2-dependent manner. Further, Snai1-KD inhibited hypoxia+TGFβ1-induced EndMT in HCAECs and FMT in HCFs *in vitro* by decreasing perinuclear/nuclear Snai1+LOXL2 expression and co-localization.

**Conclusions:** RV-specific targeting of Snai1 rescues PH-RVF by inhibiting EndMT and Fibrosis *via* a LOXL2-mediated mechanism.

## Introduction

Pulmonary hypertension (PH) is a debilitating and fatal disease that manifests initially as chronically elevated pulmonary artery (PA) pressure which leads to right ventricular (RV) hypertrophy, failure, and sudden death^1,2^. PH-induced RV failure (PH-RVF) is a significant prognostic determinant of morbidity and mortality and is characterized by cardiomyocyte hypertrophy, endothelial-to-mesenchymal transition (EndMT), fibrosis, and extracellular matrix (ECM) remodeling^3–9^. During EndMT, endothelial cells (ECs) acquire a mesenchymal cell (MC)-phenotype through a series of molecular events and develop the ability to migrate and remodel the ECM^8^. EndMT is known to contribute to cardiac fibrosis^5,10,11^. During fibrosis, fibroblasts (FBs) differentiate into functionally active myofibroblasts (MFBs), through a process known as fibroblast-to-myofibroblast transition (FMT), which can produce much more collagen compared to their fibroblast precursors^12–14^. Growing evidence from preclinical and clinical studies shows that RV fibrosis is detrimental to RV function^15–18^. Previous studies have demonstrated the close association between EndMT and fibrosis, however, the regulatory control underlying these two processes in the decompensated RV remains unclear^8^.

The EndMT-associated transcription factors mediate the endothelial-to-mesenchymal phenotypic transition^19^. The complex interplay of EndMT gatekeeper transcription factors such as Snai1 (or Snail), Snai2 (or Slug), Twist, Zeb1, and Zeb2 leads to EndMT in cancer and development^19,20^. These transcription factors interact to culminate in chromatin reorganization and their binding to cell-specific gene promoters induces EndMT^19,21–24^. In cancer and development, Snai1 recruits LOXL2, a transcriptional corepressor, to pericentromeric regions to mediate EndMT^20–22^. However, the mechanistic role of TGFβ1-Snai1-LOXL2 axis in regulating RV EndMT and fibrosis is not yet elucidated.

To address the gap in understanding the distinct molecular mechanisms of EndMT and its contribution to fibrosis in decompensated PH-RVF, we hypothesize that PH-RVF is associated with RV EndMT and fibrosis, governed by the TGFβ1-Snai1-LOXL2 axis. Furthermore, targeting Snai1 could serve as a novel therapeutic strategy for PH-RVF by reducing EndMT and fibrosis *via* a LOXL2-dependent mechanism.

In the current study, we use state-of-the-art *in vivo* and *in vitro* preclinical models and human PH-RVF data to test our hypothesis and report novel findings highlighting Snai1 as an RV-specific therapeutic target for PH-RVF.

## Materials and Methods

### Animal models of PH-RVF

All animal studies were performed in accordance with the National Institutes of Health (NIH) Guide for the Care and Use of Laboratory Animals. Protocols received UCLA animal research committee approval. Adult male Sprague Dawley rats (250–350g) received either a single subcutaneous injection of endothelial toxin Monocrotaline (MCT, 60mg/kg, MCT group, n=9) and were followed for ∼30 days or VEGF-receptor antagonist Sugen (SU5416, 20mg/kg, SuHx group, n=9) and kept in hypoxia (10% oxygen) for 3-weeks followed by 2-weeks of normoxia. Phosphate-buffered saline (PBS) treated rats served as controls (CTRL group, n=9) (Fig. S1). For *in vivo* Snai1 knockdown (Snai1-KD), MCT-rats either received Snai1-siRNA (n=7) or appropriate scramble control (siScrm: SCRM-KD; n=7) [5nM/injection every 3-4 days; 4-injections of *in vivo* ready Accell Rat SNAI1 (Custom Catalog ID: E-093687-00-0050, SMARTpool and appropriate scramble control (siScrm; Accell Non-targeting Control Pool: D-001910-01-50) commercially available from Horizon Discovery, Lafayette, CO, USA] through tail vein from day 14-30 after MCT.

### Cardiopulmonary hemodynamics

Serial non-invasive transthoracic echocardiography was performed on rats to monitor cardiopulmonary hemodynamics using a Vevo 3100 high-resolution image system (FUJIFILM VisualSonics Inc.; Toronto, Canada). Terminal echocardiography and catheterization were performed. The right and left ventricular systolic pressure (RVSP and LVSP) was measured directly by a catheter connected to a pressure transducer (ADInstruments; Colorado Springs, CO) into the RV and LV respectively.

### Morphometric analysis

The RV wall, the left ventricular (LV) wall, and the interventricular septum (IVS) were dissected and weighed. The ratio of the RV to LV plus septal weight [RV/(LV + IVS)] was calculated as the Fulton index of RV hypertrophy.

### Secondary analysis of Bulk RNA-Seq data from MCT, SuHx and PAB rat RVs

We performed secondary bioinformatics analysis and statistical analysis on the gene count matrix provided by Park et al.^23^ and Khassafi et al.^24^ on GEO using the DESeq2 R package. Genes were considered differentially expressed for an adjusted P value <0.05.

### Secondary analysis of Bulk RNA-Seq data from decompensated Human PH-RVF

RNA-seq data from human control and decompensated PAH-RVF (dRVs) were obtained from NCBI GEO database GSE198618^9^. We performed secondary bioinformatics and statistical analysis on the gene count matrix using the DESeq2 R package. Genes were considered differentially expressed for an adjusted P value <0.05.

### RNA isolation and qPCR

**Rat:** Total RNA was isolated from rat RV and lung tissue using Trizol (Invitrogen). Quantitative real-time PCR was performed following the protocol for the “mixed primer strategy” (Random primer + Oligo dT) using the SsoAdvanced Universal SYBR Green Supermix (BioRad, Cat #172-5274) in the CFX Connect™ Real-Time PCR Detection System (384-well block, Biorad, Cat #1855201). GADPH was used as an internal reference control, and relative gene expression was normalized to the PBS-treated group (primer details as described in Table S1).

**Human:** Total RNA was extracted from human RV tissue using TRIzol (Ambion by Life Technologies, Ref#15596018). cDNA was prepared using qScript Flex cDNA Synthesis Kit from (Quanta Bio, Cat #95049-100). Quantitative real-time PCR was performed following the protocol for the “mixed primer strategy” (Random primer + Oligo dT) using the SsoAdvanced Universal SYBR Green Supermix (BioRad, Cat #172-5274) in the QuantStudio™ 7 Flex Real-Time PCR System (384-well block, Applied Biosystems, Cat #4485701). 18s rRNA was used as an internal reference control, and relative gene expression was normalized to the control group (primer details as described in Table S2).

### Western blot analysis

**Rat:** Protein lysates were prepared from rat RV tissue using modified RIPA lysis buffer containing protease and phosphatase inhibitor cocktails (RIPA: Cat# R0278, PMSF: Cat# P7626, Protease Inhibitor Cocktail: Cat# P3840, and Phosphatase Inhibitor Cocktail 2: P5726, Sigma). Protein estimation was performed using DC protein assay (DC™ Protein Assay Kit II, Cat# 5000112, Biorad). Proteins were diluted in 4x Laemmli sample buffer, boiled, separated on 4-20% gradient gels (Bio-Rad) by SDS page and subsequently transferred onto nitrocellulose membranes (Bio-Rad) using semi-dry blotting (Trans-Blot® Turbo™ Transfer System, Bio-Rad, #1704150). After transfer, membranes were blocked with 5% bovine serum albumin (Sigma) and incubated overnight at 4°C with primary antibodies directed against Snai1 (CST#3879; 1:1000), LOXL2 (NBP1-32954; 1:1000), GAPDH (CST14C10#2118; 1:5000) and Vinculin (Sigma #V9131; 1:5000) (as described in Table S3). The blots were incubated with IR Dye-conjugated secondary antibodies (as described in Table S4) for 2 h at room temperature, and visualized, scanned and quantified using the LI-COR Odyssey™ Infrared Imaging System and Image Studio Lite.

**Human:** Protein lysates were prepared from human RV tissue using modified RIPA lysis buffer containing protease and phosphatase inhibitor cocktails (Sigma). Proteins were diluted in 4x Laemmli sample buffer, boiled, loaded (10ug of protein per loading), and separated on mini-PROTEAN TGX (4-20%) (Bio-Rad, Cat #4561094), and subsequently transferred onto PVDF membranes (Bio-Rad) using semi-dry blotting (Trans-Blot® Turbo™ Transfer System, Bio-Rad, #1704150). After transfer, membranes were blocked with 5% nonfat-dried milk bovine (Sigma, M7409-5BTL) and incubated overnight at 4°C with primary antibodies directed against Snai1 (CST#3879; 1:1000), LOXL2 (NBP1-32954; 1:1000) in TBS-T, and Amidoblack. IR Dye-conjugated secondary antibodies (LI-COR) were used for detection and blots were scanned and quantified using the ChemiDoc MP Imaging System (Biorad, #12003154) and software.

### Histology and Trichrome staining

Transverse 4- to 6-um sections were obtained from cryoblocks of embedded RV and Lungs with a cryostat. Paraffin-embedded control and PH-RVF human RV sections, obtained from the UCLA pathology laboratory, were sectioned at 5um. Masson’s trichrome staining was performed according to the manufacturer’s protocol, and images were acquired using a confocal microscope (Nikon).

### Immunofluorescence and Confocal Microscopy

Before collecting patient RV samples, we obtained institutional review board approval from the Office of the Human Research Protection Program at the University of California, Los Angeles, and informed consent from all subjects, and confirmed that the intended experiments conformed to the principles set out in the WMA Declaration of Helsinki and the Department of Health and Human Services Belmont Report. Paraffin-embedded human RV sections were rehydrated, and antigen retrieval was performed by heat-induced epitope retrieval at pH 6. Tissue sections were blocked for an hour at room temperature in PBS with 5% goat serum, 1% BSA, 0.1% Triton, 0.05% Tween, and 0.3 M glycine. Primary antibodies were incubated overnight at 4°C and diluted as described in Table S3 in PBS with 1% BSA and 0.5% Triton. Secondary antibodies were incubated for an hour at room temperature and diluted as described in Table S4 in PBS with 0.05% Tween. All images were acquired using a confocal microscope (Nikon).

### *In vitro* experiments

**HCAECs:** Human coronary artery endothelial cells (HCAECs: Promocell# C-12221) were incubated in hypoxia condition in the hypoxia chamber with TGFβ1 (recombinant human TGFβ1: 15ng/ml, Cat #T7039, Sigma)^25^ for 72 h. Snai1-KD was performed in HCAECs under hypoxia and TGFβ1 (15ng/ml) treatment using Human SNAI1 siRNA [1uM/ml of medium of *in vitro* ready Accell Human SNAI1siRNA-SMARTpool (6615), and appropriate scramble control (siScrm; D-001910-01-50) commercially available from Horizon Discovery, Lafayette, CO, USA] in combination with the Lipofectamine™ RNAiMAX Transfection Reagent (Cat# 13778100, Invitrogen) for enhanced transfection efficiency, following the manufacturer’s instructions.

**qPCR:** Total RNA was isolated from the HCAECs using RNeasy Mini Kit (Cat #74104 and 74106, QIAGEN). Quantitative real-time PCR was performed following the protocol for the “mixed primer strategy” (Random primer + Oligo dT) using the SsoAdvanced Universal SYBR Green Supermix (BioRad, Cat #172-5274) in the CFX Connect™ Real-Time PCR Detection System (384-well block, Biorad, Cat #1855201). RPLP0 was used as an internal reference control, and relative gene expression was normalized to the PBS- or Scrm-treated group (primer details as described in Table S5).

### Western blot analysis

Cells were lysed using modified RIPA lysis buffer containing protease and phosphatase inhibitor cocktails (RIPA: Cat# R0278, PMSF: Cat# P7626, Protease Inhibitor Cocktail: Cat# P3840, and Phosphatase Inhibitor Cocktail 2: P5726, Sigma). Protein estimation was performed using DC protein assay (DC™ Protein Assay Kit II, Cat# 5000112, Biorad). Proteins were diluted in 4x Laemmli sample buffer, boiled, separated on 4-20% gradient gels (Bio-Rad) by SDS page and subsequently transferred onto nitrocellulose membranes (Bio-Rad) using semi-dry blotting (Trans-Blot® Turbo™ Transfer System, Bio-Rad, #1704150). After transfer, membranes were blocked with 5% bovine serum albumin (Sigma) and incubated overnight at 4°C with primary antibodies directed against Snai1 (CST#3879; 1:1000), LOXL2 (NBP1-32954; 1:1000), GAPDH (CST14C10#2118; 1:5000) (as described in Table S3). The blots were incubated with diluted IR Dye-conjugated secondary antibodies (as described in Table S4) for 2 h at room temperature, and visualized, scanned and quantified using the LI-COR Odyssey™ Infrared Imaging System and Image Studio Lite.

**Immunofluorescence:** HCAECs were fixed in 4% PFA, washed with PBS, and incubated for an hour with the blocking solution, primary antibodies, and secondary antibodies (as described in Table S3-4). All images were acquired using a confocal microscope (Nikon).

**HCFs:** Human Cardiac Fibroblasts (HCFs: Promocell# C-12375) were incubated in hypoxia condition in the hypoxia chamber with TGFβ1 (recombinant human TGFβ1: 15ng/ml, Cat # T7039, Sigma)^10^ for 72 h. Snai1-KD was performed in HCFs under hypoxia and TGFβ1 (15ng/ml) treatment using Human SNAI1 siRNA (Accell siRNA-SNAI1 SMARTpool, *in vitro* ready, Horizon Discovery; 1uM/ml of medium) in combination with the Lipofectamine™ RNAiMAX Transfection Reagent (Cat# 13778100, Invitrogen) for enhanced transfection efficiency, following the manufacturer’s instructions.

**qPCR:** We followed the same protocol as mentioned before under the qPCR subsection for the HCAECs.

**Western blot analysis:** We followed the same protocol as mentioned before under the qPCR subsection for the HCAECs.

**Immunofluorescence:** HCFs were fixed in 4% PFA, washed with PBS, and incubated for an hour with the blocking solution, primary antibodies, and secondary antibodies as described above. All images were acquired using a confocal microscope (Nikon).

### Patient Demographics

Patient demographics of the human samples used in this study are described in Table S6 (Table. S6).

### Statistical analysis

Data are presented as mean ± SEM. Comparisons between 2 groups were performed using unpaired Student’s t-test and 3 groups were performed using one-way ANOVA (with Tukey or Dunnett post-hoc tests) using GraphPad Prism 9 Software (San Diego, CA, USA). Bioinformatic analyses of omics data were performed using R version 4.

## Results

### Development of severe PH-induced RVF in MCT and SuHx Rats

We utilized well-established rat models of severe PH-induced RVF induced by MCT and SuHx. Severe PH and RVF were confirmed using serial transthoracic echocardiography and terminal right heart catheterization (Fig. 1A-H). Both MCT and SuHx rats developed significant PH as evidenced by significantly increased RV systolic pressure (RVSP) (MCT=82.32±2.17; SuHx=72.46±5.03, vs. control=26.44±0.85 mmHg; p<0.0001 MCT vs. control, p<0.0001 SuHx vs. control; Fig. 1C). RV dysfunction was demonstrated by increased RV internal diameter at end-diastole (RVIDd) (MCT=3.42±0.21; SuHx=3.41±0.32 vs. control=1.29±0.08 mm; p<0.0001 MCT vs. control, p<0.0001 SuHx vs. control; Fig. 1D) and decreased RV fractional area change (RVFAC%) (MCT=15.09±1.10; SuHx=15.15±1.61 vs. control=35.29±1.36%; p<0.0001 MCT vs. control, p<0.0001 SuHx vs. control; Fig. 1E) in MCT and SuHx rats. MCT and SuHx rats also demonstrated significantly increased RV hypertrophy evaluated as Fulton index (RV/LV+IVS) (MCT=0.75±0.05; SuHx=0.81±0.06 vs. control=0.29±0.01; p<0.0001 MCT vs. control, p<0.0001 SuHx vs. control; Fig. 1F). No significant differences were documented between SuHx and MCT rats in all of the above-mentioned parameters. Further, no significant change was observed in left ventricular ejection fraction (LVEF%) (MCT=78.40±1.33; SuHx=80.28±3.94 vs. control=76.36±2.71; p= 0.5582 MCT vs. control, p=0.5880 SuHx vs. control; Fig. 1G) and left ventricular systolic pressure (LVSP) (MCT=84.79±4.87; SuHx=98.35±5.25 vs. control=81.66±2.89 mmHg; p=0.8656 MCT vs. control, p=0.0879 SuHx vs. control; Fig. 1H) in MCT and SuHx rats compared to control (Fig. 1G-H).

**Fig. 1.**
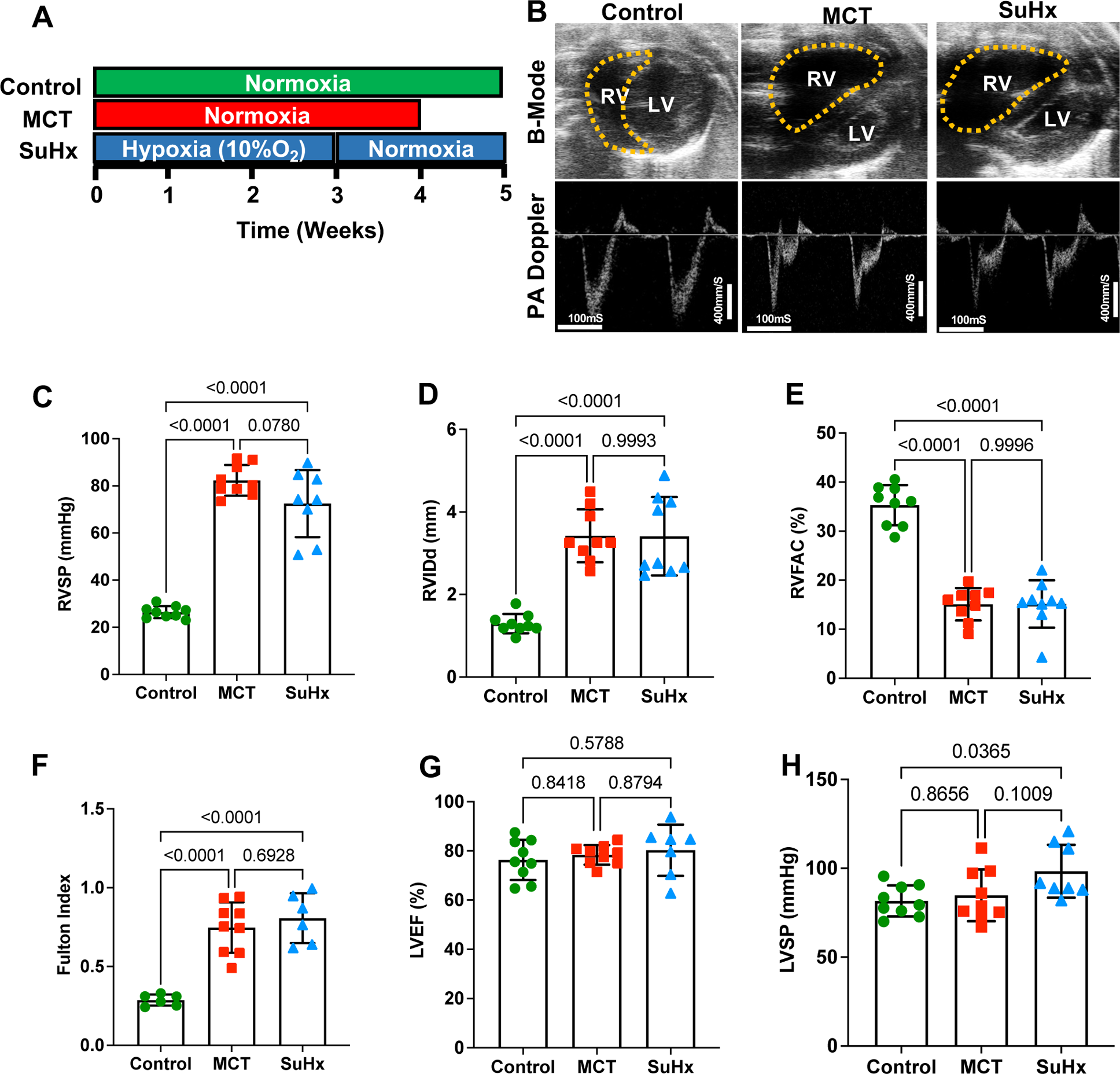
Development of RV failure in MCT and SuHx PH-RVF rats. (**A**) Experimental protocol for MCT, SuHx, and Ctrl rats. (**B**) Echocardiographic B-mode images at end-diastole and PA Doppler images from parasternal short-axis views. (**C**) RVSP, (**D-E**) a summary of echocardiographic assessment of RV function (RVID_d_, RVFAC), (**F**) Fulton index (RV hypertrophy), (**G**) LVEF and (**H**) LVSP. n=6-9 per group. RVSP: RV systolic pressure, RVID_d_: rRV internal diameter at end-diastole, RVFAC: RV fractional area change, LVEF: LV ejection fraction, LVSP: LV systolic pressure.

### Significant concordance in genes and pathways related to EndMT and fibrosis between rat and human decompensated RVs

Secondary bioinformatic analysis of RV-RNA-Seq data from the dRV of MCT, SuHx, and human PH-RVF highlighted a significant overlap of genes (987 genes) and pathways (23 pathways) between all 3 datasets (Fig. 2A). Interestingly, out of these 23 common pathways, EndMT was the top common upregulated pathway in the failing RV of both MCT and SuHx rat models and PH-RVF patients (Fig. 2A)^9,23^. Several key driver genes underlying EndMT, and fibrosis were found to be commonly upregulated in all the 3 groups compared to their respective controls. Further, secondary analysis of RV-RNA-Seq data from the dRV of MCT, SuHx, PAB, and human PH-RVF highlighted a significant overlap of genes (450 genes) and pathways (11 pathways) between all 4 datasets (Fig. 2B). Interestingly, out of these 11 common pathways, EMT/EndMT was again found to be the top common upregulated pathway in the failing RV of MCT, SuHx, PAB rat models, and PH-RVF patients (Fig. 2B)^9,23,24^. In addition, pathway enrichment analysis also demonstrated TGFβ signaling, and hypoxia as the other highly upregulated EMT/EndMT, FMT, and fibrosis-related pathways in the decompensated RV of MCT SuHx, and PAB rats and human PH-RVF (Fig. 2E-F). Several key driver genes underlying EndMT, FMT, and fibrosis were found to be commonly upregulated in all 4 groups compared to their respective controls (Fig. 2C-D). From the common EndMT-inducing transcription factors (Snai1/Snail, Snai2/Slug, Twist1/2, Zeb1/2), only Snai1 was preferentially upregulated in dRV of MCT, SuHx, and PAB rats and was also found to be significantly increased in the dRV of patients with PH-RVF (Fig. S2D-E). Interestingly, Snai1 and LOXL2 were upregulated in the compensated RV of MCT and PAB rats and were also found to be significantly increased in the compensated RV of patients with PH-RVF (Fig. S2A).

**Fig. 2.**
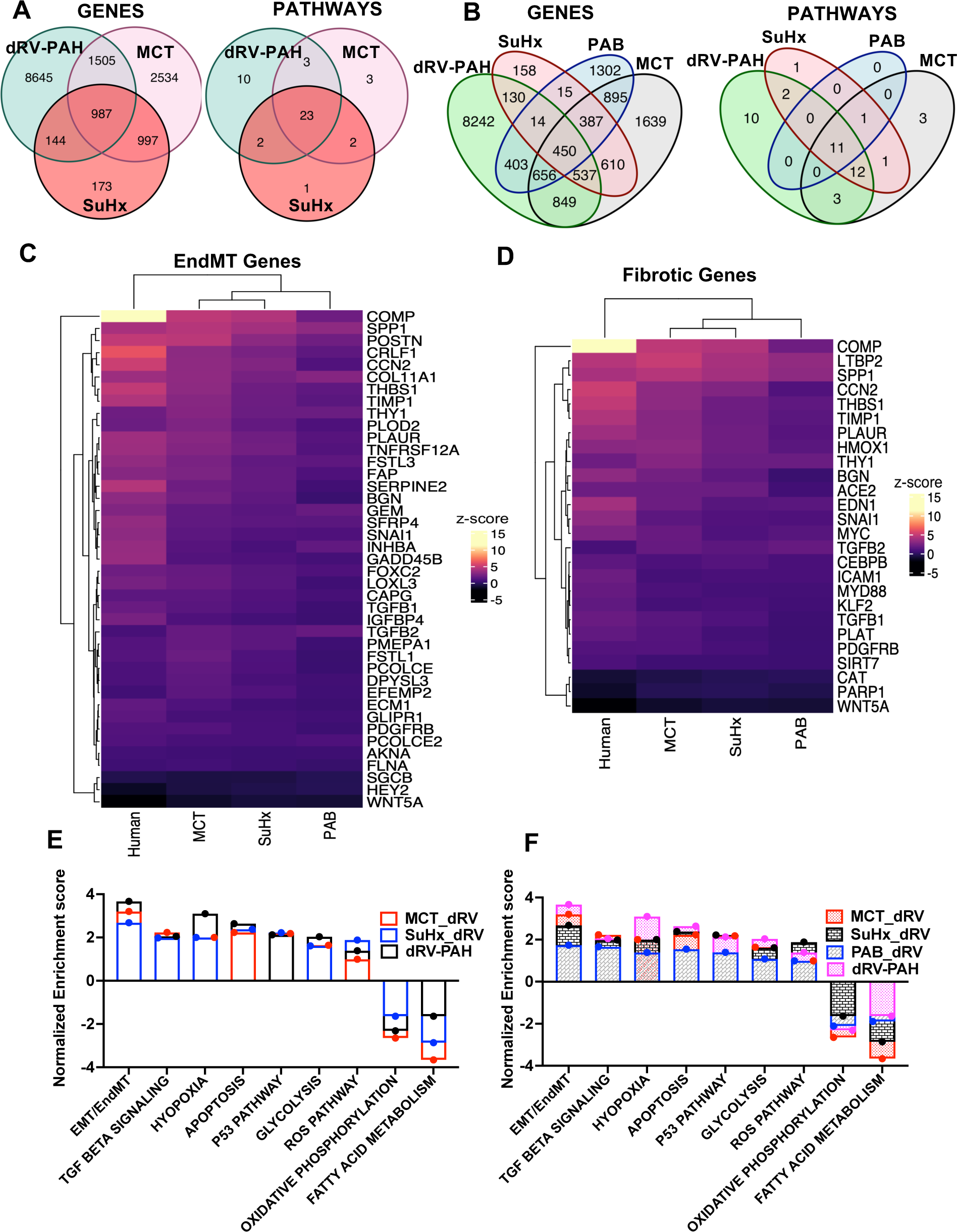
Secondary analysis of RNASeq data from the decompensated RV of MCT, SuHx, and PAB rats, and human PH-RVF. **(A)** Venn diagrams showing the total number and overlapping genes and pathways in the decompensated RVs (dRVs) of MCT, SuHx rats, and human PH-RVF. **(B)** Venn diagrams showing the total number and overlapping genes and pathways in the dRVs of MCT, SuHx, PAB rats, and human PH-RVF. **(C-D)** Heatmaps of EndMT and Fibrotic genes showing the concordance in their differential expression pattern in the dRVs of MCT, SuHx, PAB rats, and human PH-RVF **(E)** Hallmark pathway enrichment analysis of overlapping pathways in the dRVs of MCT, SuHx rats and human PH-RVF. **(F)** Hallmark pathway enrichment analysis of overlapping pathways in the dRVs of MCT, SuHx, PAB rats, and human PH-RVF.

### Establishment of Snai1 and related network (TGFβ1-Snai1-LOXL2 axis) as the key regulatory hub underlying RV

As TGFβ1 is known to upregulate Snai1^10,26,27^, which recruits LOXL2 to induce EMT/EndMT^28^, we assessed the expression of TGFβ1-Snai1-LOXL2 axis in dRVs from MCT, SuHx, and PAB rat models and human PH-RVF. Interestingly, from our RV RNA seq data analysis, we found TGFβ1-Snai1-LOXL2 axis was significantly upregulated in dRVs from MCT, SuHx, and PAB rat models and human PH-RVF (Fig. S2B-C).

### Validation of Snai1 and its association with LOXL2 in RV EndMT in two severe rat models and patients with PH-RVF

Significantly increased expression of Snai1 was associated with LOXL2 and α-SMA upregulation in the vascular ECs undergoing EndMT in the RV of MCT and SuHx rats compared to control (Fig. 3A, Fig. S3). Further, significantly increased TGFβ1, Snai1, and LOXL2 at both transcripts (TGFβ1: MCT vs control, p=0.0008, SuHx vs control, p=0.0064, MCT vs SuHx, p=0.2632; Snai1: MCT vs control, p=0.0162, SuHx vs control, p=0.0178, MCT vs SuHx, p=0.8724; LOXL2: MCT vs control, p=0.0008, SuHx vs control, p=0.0172; MCT vs SuHx, p=0.1034, Fig. 3C-E) and protein levels were documented in the RV of MCT and SuHx rats (Snai1: MCT vs control, p=0.033, SuHx vs control, p=0.0383, MCT vs SuHx, p=0.6754; LOXL2: MCT vs control, p=0.0007, SuHx vs control, p=0.0004; MCT vs SuHx, p=0.0462, Fig. 3B).

**Fig. 3.**
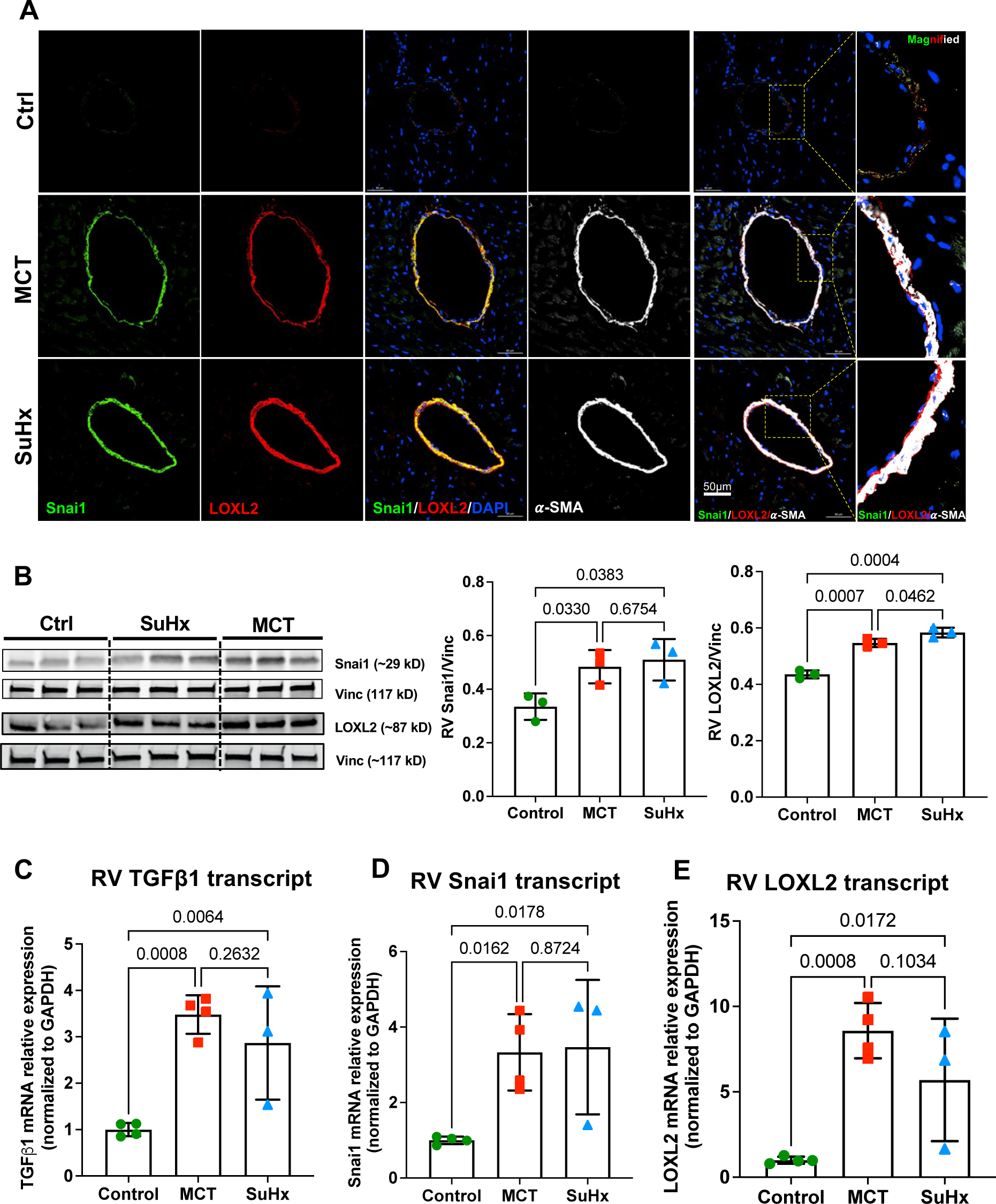
Validation of EndMT in the RV of MCT and SuHx rats with PH-RVF. **(A)** Triple immunolabeling of Snai1, LOXL2, and α-SMA (mesenchymal marker) in the dRVs demonstrated increased EndMT with increased colocalization of Snai1 and LOXL2 (their association with EndMT) in the RVs of SuHx and MCT rats. (**B)** WB and densitometric analysis demonstrated significantly increased Snai1 and LOXL2 in the RVs of SuHx and MCT rats. (**C-E)** qPCR analysis demonstrated significantly increased expression of TGFβ1, Snai1, and LOXL2 transcripts in the RVs of SuHx and MCT rats. N=3-4 per group.

Similarly, significantly increased Snai1 and LOXL2 proteins (SNAI1: PAH vs ctrl, p=0.0288, LOXL2: p=0.0068, Fig. 4A) and transcripts (SNAI1: PAH vs ctrl, p=0.0225; LOXL2: p=0.0156, Fig. 4B) were documented in the RV of patients with PH-RVF, validating the significantly increased Snai1 and LOXL2 gene expression in PH-RVF patient RV-RNAseq data (Snai1: PAH vs ctrl, p=0.0162, SuHx vs control, p=0.0178, MCT vs SuHx, p=0.8724; LOXL2: MCT vs control, p=0.0008, SuHx vs control, p=0.0172; MCT vs SuHx, p=0.1034, Fig. 4A-B). Further, in the RV of patients with PH-RVF, Snai1 was increased and colocalized with α-SMA and vWF in the IPAH and SSC-associated PH-RVF showing the association of Snai1 with vascular EndMT (Fig. 4C). Similarly, increased Snai1 was associated with LOXL2 upregulation, and co-localized with LOXL2 in the vasculature (Fig. 4D). Interestingly, increased Snai1 and LOXL2 immunoreactivity along with their co-localization was documented in the nuclear/perinuclear regions in the interstitial space of the RV of patients with PH-RVF (Fig. 4E).

**Fig. 4.**
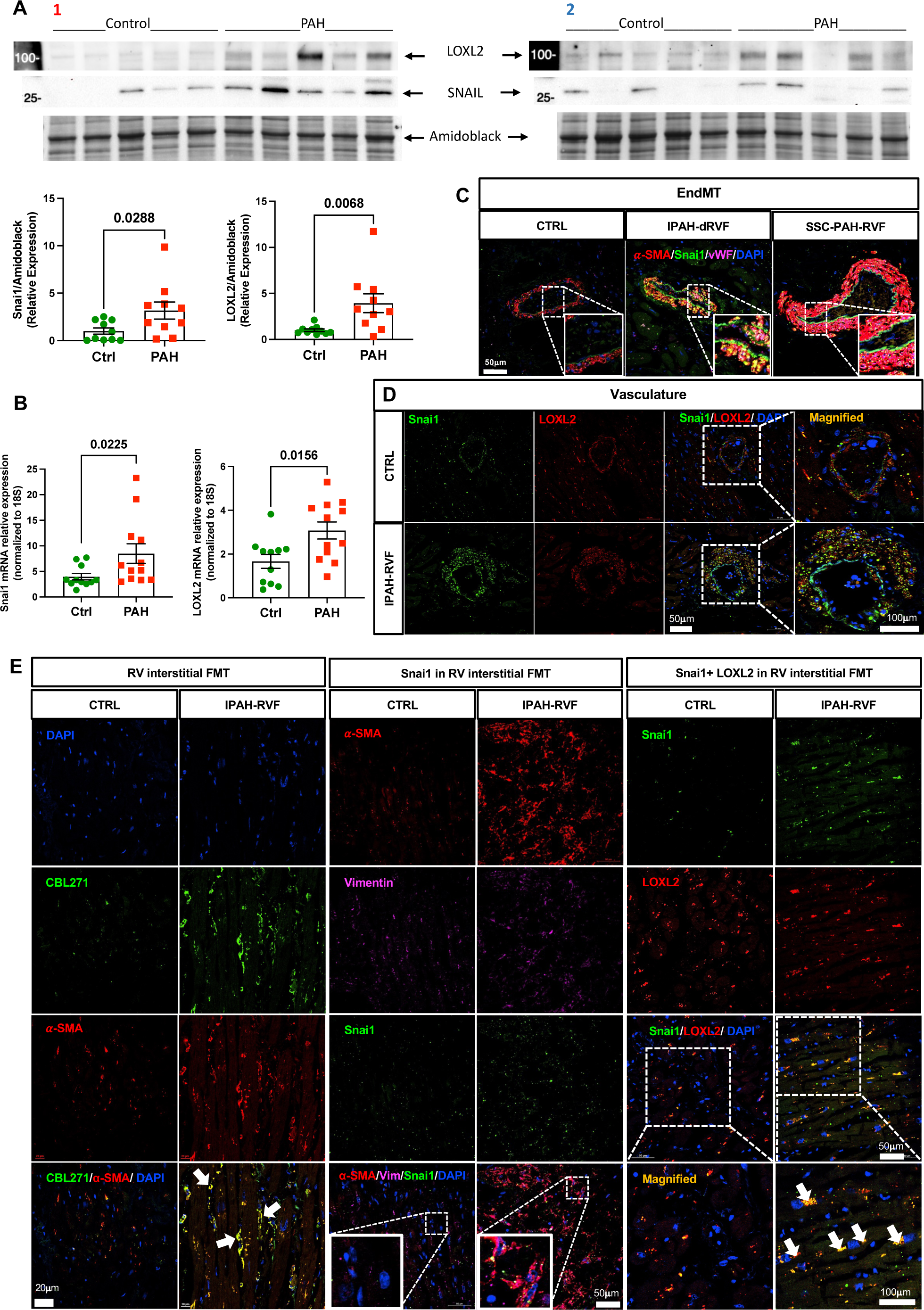
Validation of Human RV EndMT and FMT. **(A)** WB and densitometric analysis demonstrated significantly increased Snai1 and LOXL2 in the RVs of human PH-RVF from two different patient cohorts (cohort1 and 2). N=10 per group. **(B)** qPCR analysis demonstrated significantly increased expression of Snai1 and LOXL2 transcripts in the RVs of human PH-RVF. N=10 per group. **(C)** Representative immunofluorescence images with α-SMA (red), VWF (magenta), Snai1 (green), and DAPI (blue) in human RV showed significantly increased EndMT associated with increased Snai1. **(D)** Representative immunofluorescence images with Snai1 (green), LOXL2 (red), and DAPI (blue) in human RV demonstrated significantly increased colocalization of Snai1 and LOXL2 in the vasculature. **(E)** In the left panel, representative immunofluorescence images with CBL271 (green), α-SMA (red), and DAPI (blue) demonstrated significantly increased FMT in human RV interstitial space. In the middle panel, representative triple labeling immunofluorescence images with α-SMA (red), Vimentin (magenta), Snai1 (green), and DAPI (blue) demonstrated that significantly increased Snai1 is associated with increased interstitial FMT in human RV. In the right panel, representative immunofluorescence images with Snai1 (green), LOXL2 (red), and DAPI (blue) showed increased immunoreactivity and colocalization of Snai1 and LOXL2 in the RV interstitial space of the human PH-RVF. N=3 per group.

### Hypoxia and TGFβ1 induce EndMT in HCAECs and FB-to-MFB transition in HCFs *via* a Snai1-LOXL2-dependent mechanism

In addition to EndMT, hypoxia, and TGFβ1 signaling were significantly upregulated hallmark pathways in the dRV of MCT, SuHx, and PAB rat models and human PH-RVF (Fig. 2E-F). Hence, to mimic or recapitulate the *in vivo* rat and human pathophysiological conditions *in vitro* in our cell culture models, next, we decided to treat HCAECs and HCFs with Hypoxia and TGFβ1 in combination to check if the combined hypoxia and TGFβ1 stimulus synergistically induces EndMT in HCAECs and FB-to-MFB transition in HCFs (Fig. S4). We also checked whether EndMT in HCAECs and FB-to-MFB transition in HCFs are mediated *via* the Snai1-LOXL2-dependent mechanism. Interestingly, hypoxia+TGFβ1 treatment induced EndMT in the HCAECs as evidenced by double immunolabeling with endothelial marker vWF and mesenchymal marker a-SMA (Fig. 5A). EndMT resulted in increased nuclear/peri-nuclear co-localization of Snai1+LOXL2, suggesting a possible involvement of the TGFβ1, Snai1, and LOXL2 in EndMT (Fig. 5B). Similarly, hypoxia+TGFβ1 treatment induced FB-to-MFB transition in the HCFs as documented by double immunolabeling with FB marker Vimentin and MFB marker a-SMA. FB-to-MFB transition resulted in increased nuclear/peri-nuclear co-localization of Snai1+LOXL2, indicating a possible role of the TGFβ1, Snai1, and LOXL2 in FB-to-MFB transition (Fig. 5C-D).

**Fig. 5.**
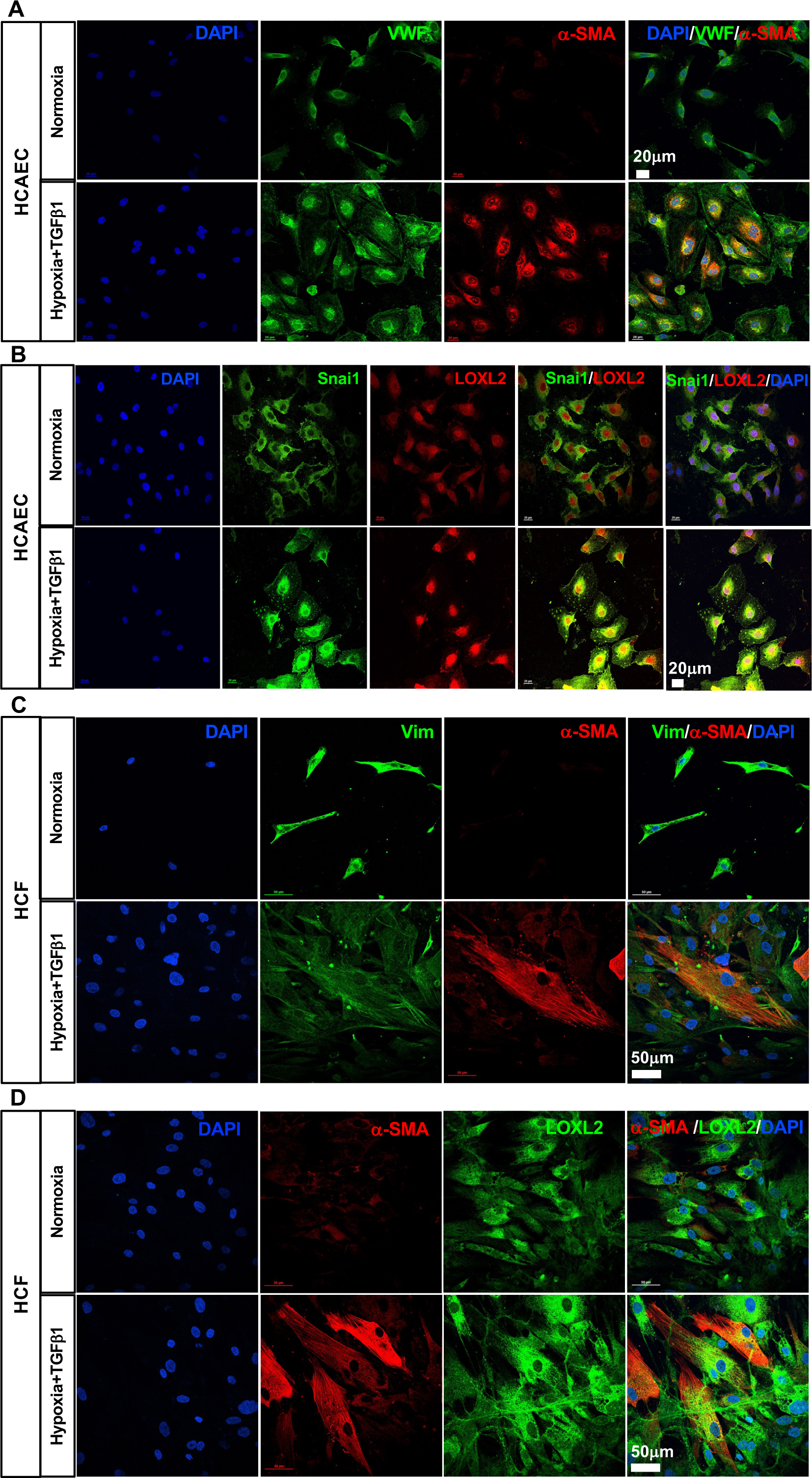
Hypoxia and TGFβ1 induce EndMT in HCAECs and FMT in HCFs *in vitro*. **(A)** Representative immunofluorescence images with VWF (green), α-SMA (red), and DAPI (blue) demonstrated induction of EndMT in HCAECs *in vitro* under Hypoxia and TGFβ1 treatment for 72 hrs**. (B)** Representative immunofluorescence images with Snai1 (green), LOXL2 (red), and DAPI (blue) demonstrated intense perinuclear/nuclear immunoreactivity and colocalization of Snai1 and LOXL2, indicating the involvement of Snai1 and LOXL2 in inducing EndMT in HCAECs *in vitro* under Hypoxia and TGFβ1 treatment for 72 hrs. **(C)** Representative immunofluorescence images with Vimentin (green), α-SMA (red), and DAPI (blue) demonstrated induction of FMT in HCFs under Hypoxia and TGFβ1 treatment for 72 hrs**. (D)** Representative immunofluorescence images with α-SMA (red), LOXL2 (green), and DAPI (blue) demonstrated intense perinuclear/nuclear immunoreactivity of LOXL2, suggesting the involvement of LOXL2 in inducing FMT in HCFs under Hypoxia and TGFβ1 treatment for 72 hrs. N=3 per group.

### Snai1 Knockdown rescues PH-induced RV Failure in MCT rats by inhibiting RV EndMT, FB-to-MFB transition, and fibrosis via a LOXL2-dependent mechanism

Snai1-KD resulted in significantly reduced RVSP (si-Snai1=46.31±1.83; vs. si-Scrm=73.68±6.26 mmHg; p=0.0012 si-Snai1 vs. si-Scrm; Fig. 6A-B), decreased RV hypertrophy (Fulton index) (si-Snai1=0.43±0.02; vs. si-Scrm=0.74±0.05 mmHg; p<0.0001 si-Snai1 vs. si-Scrm; Fig. 6C) and RVIDd (si-Snai1=1.43±0.05; vs. si-Scrm=3.01±0.42 mmHg; p= 0.0059 si-Snai1 vs. si-Scrm; Fig. 6D) in MCT rats with Snai1-KD compared to scramble-treated MCT rats. Snai1-KD significantly improved RVFAC% (si-Snai1=24.65±1.05; vs. si-Scrm=11.12±0.56 mmHg; p<0.0001 si-Snai1 vs. si-Scrm; Fig. 6E) in MCT rats with Snai1-KD compared to scramble-treated MCT rats. No significant differences were documented in the LVEF (si-Snai1=76.58±2.46; vs. si-Scrm=77.56±3.55 mmHg; p=0.8301 si-Snai1 vs. si-Scrm; Fig. 6F) and LVSP (si-Snai1=94.21±5.71; vs. si-Scrm=95.13±6.52 mmHg; p=0.9191 si-Snai1 vs. si-Scrm; Fig. 6G) in MCT rats with Snai1-KD compared to scramble-treated MCT rats.

**Fig. 6.**
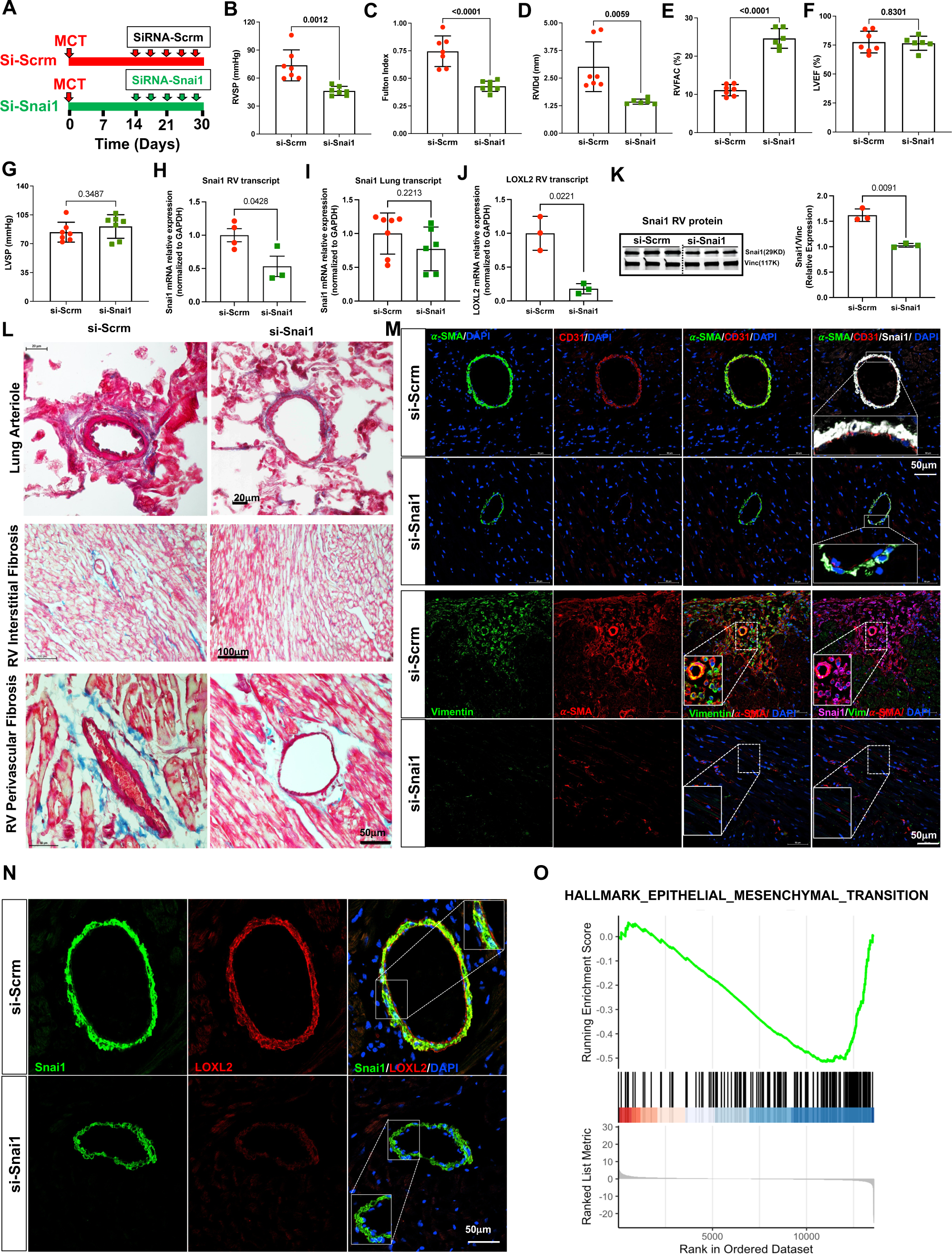
Snai1 KD rescues MCT PH-induced RV failure. **(A)** Experimental protocol for *in vivo* Snai1 knockdown, MCT-rats either received Snai1-siRNA or Scramble from day-14 to day-30 after MCT, **(B-G)** RVSP, Fulton Index, RVIDd, RVFAC, LVEF, LVSP (N=6-7 per group). **(H)** RV Snai1 transcript, **(I)** Lung Snai1 transcript, **(J)** RV LOXL2 transcript, **(K)** Snai1 protein in RV by WB, **(L)** Trichrome staining of lung cross-sections demonstrating arteriolar thickness of pulmonary arterioles, and RV transverse-sections demonstrating interstitial and peri-vascular fibrosis (blue), **(M)** In the top panel, representative immunofluorescence images with α-SMA (green), CD31 (red), Snai1 (white), and DAPI (blue) assessing EndMT and Snai1 expression in RV vasculature. In the bottom panel, representative immunofluorescence images with Vimentin (green), α-SMA (red), Snai1 (magenta), and DAPI (blue) assessing FMT and Snai1 expression in RV interstitial space**. (N)** Representative immunofluorescence images with Snai1 (green), LOXL2 (red), and DAPI (blue) assessing the expression and colocalization of Snai1 and LOXL2 in the vasculature of RV sections. **(O)** RV RNASeq Hallmark pathway-specific Gene-set Enrichment analysis Pathway Enrichment analysis showing Snai1-KD mediated successful reversal of RV EMT/EndMT in PH-RVF MCT rats. N=3-4 per group. FDR<0.05.

We confirmed the successful KD of Snai1 in the RV using qPCR (p=0.0482; Fig. 6H) and WB (p=0.0091; Fig. 6K). Interestingly, Snai1-KD resulted in a significant reduction in LOXL2 expression in the RV, suggesting a LOXL2-mediated rescue of RVF by Snai1-KD (p=0.0221; Fig. 6J). However, no change was observed in lung Snai1 transcript (p=0.2213; Fig. 6I) levels between these two groups.

Further, Trichrome staining revealed significantly reduced lung vascular (arteriolar) wall thickness (Fig. 6L, middle panel) in the Snai1-KD MCT rats compared to scramble-treated MCT rats. Trichrome staining also demonstrated significantly reduced RV perivascular and interstitial fibrosis in Snai1-KD MCT rats (Fig. 6L, bottom panel). Our exciting results demonstrated that Snai1-KD results in the rescue of PH, RV hypertrophy, and RV failure in MCT rats. Significantly reduced Snai1 expression and its association/colocalization with vascular EndMT and interstitial FB-to-MFB transition were also observed in Snai1-KD MCT rats compared to the scramble-treated MCT rats using markers of EndMT (CD31 and α-SMA) and FMT (Vim and α-SMA). Findings demonstrated that Snai1-KD inhibits RV EndMT, FMT, and fibrosis (Fig. 6M). Further investigations showed increased expression and colocalization of Snai1 and LOXL2 in the vasculature (mostly in vascular ECs) (Fig. 6N) and increased expression of Snai1 in the interstitial space (mostly in interstitial FBs or MFBs) (Fig. 6M). Next, we processed the RVs from both Snai1-KD and scramble-treated MCT rats for bulk RNA sequencing and found EMT/EndMT as the top-downregulated pathway. In addition to EMT/EndMT (Fig. 6O), hypoxia, TGFβ signaling, and apoptosis (Fig. S5) were the significantly downregulated hallmark pathways after Snai1-KD in MCT rats. All the key EndMT, FMT, and fibrosis driver genes (Fig. S5) commonly upregulated in the decompensated RV of MCT, SuHx, and PAB rats, and human PH-RVF compared to their respective controls, were significantly down-regulated in the rescued RV of Snai1-KD MCT rats.

### Snai1 KD Inhibits Hypoxia+TGFβ1-induced EndMT in HCAECs and FB-to-MFB Transition in HCFs via reducing LOXL2 *In Vitro*

Snai1-KD in HCAECs under Hypoxia+TGFβ1 *in vitro* (Fig. 7A, S6) resulted in significant reduction in Snai1 expression (p=0.0127; Fig. 7B) as demonstrated by qPCR analysis. Snai1-KD also resulted in a significant reduction of LOXL2 (p=0.0286; Fig. 7C), and α-SMA expression (p=0.0272; Fig. 7D) in HCAECs under Hypoxia+TGFβ1 *in vitro*. Snai1-KD inhibited EndMT as documented by reduced α-SMA in HCAECs by decreasing Snai1 and LOXL2 co-localization in the nuclear and perinuclear areas (Fig. 7E). Immunoblots and corresponding densitometric analysis also showed significantly reduced Snai1 (p=0.0032; Fig. 7F), LOXL2 (p=0.0031; Fig. 7G) and α-SMA (p=0.0166; Fig. 7H) in the Snai1-KD HCAECs and further validated our immunofluorescence findings.

**Fig. 7.**
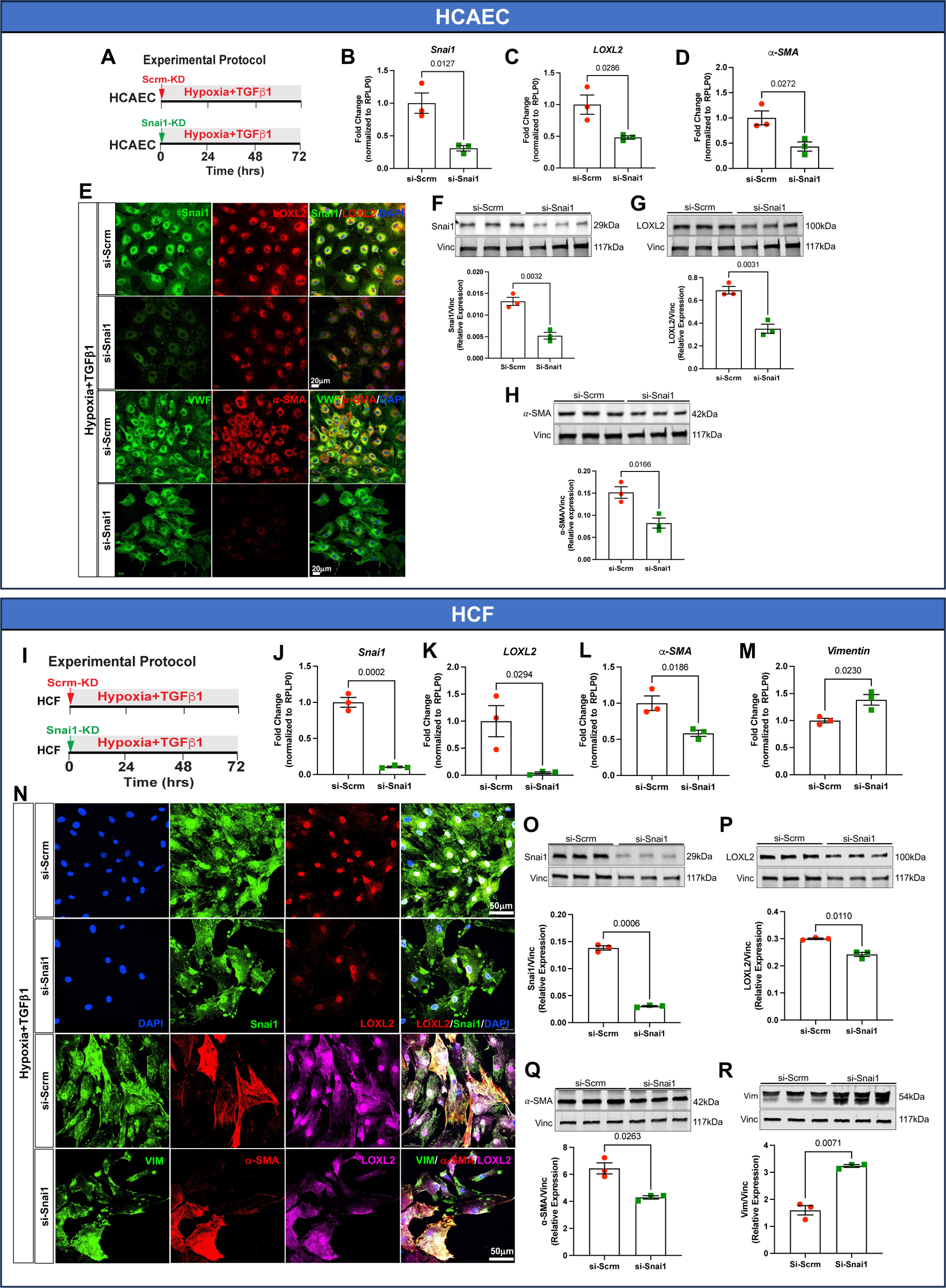
Snai1-KD inhibits EndMT in HCAEC and FMT in HCF under hypoxia and TGFβ1 *in vitro via* LOXL2-mediated mechanism. **(A)** Experimental Protocol. **(B-D)** Confirmation of successful Snai1-KD by siRNA and a significant reduction in LOXL2 and α-SMA in HCAEC by qPCR (n=3 per treatment group). **(E)** In the top panel, representative immunofluorescence images with Snai1 (green), LOXL2 (red), and DAPI (blue) assessing Snai1 and LOXL2 expression in HCAEC under hypoxia and TGFβ1 with scramble or Snai1 siRNA. In the bottom panel, representative immunofluorescence images with vWF (green), α-SMA (red), and DAPI (blue) assessing EndMT in HCAEC under hypoxia and TGFβ1 with scramble or Snai1 siRNA**. (F-H)** Confirmation and validation of successful Snai1-KD by siRNA and a significant reduction in LOXL2 and α-SMA in HCAEC by WB and corresponding densitometric analysis (n=3 per treatment group). Snai1-KD inhibits hypoxia+TGFβ1-induced EndMT by reducing Snai1 and LOXL2 expression and colocalization. **(I)** Experimental Protocol. **(J-M)** Confirmation of successful Snai1-KD by siRNA and a significant reduction in LOXL2 and α-SMA in HCF by qPCR (n=3 per treatment group). **(N)** In the top panel, representative immunofluorescence images with Snai1 (green), LOXL2 (red), and DAPI (blue) assessing Snai1 and LOXL2 expression in HCF under hypoxia and TGFβ1 with scramble or Snai1 siRNA. In the bottom panel, representative immunofluorescence images with vWF (green), α-SMA (red), and DAPI (blue) assessing FMT in HCF under hypoxia and TGFβ1 with scramble or Snai1 siRNA**. (O-R)** Confirmation and validation of successful Snai1-KD by siRNA and a significant reduction in LOXL2 and α-SMA in HCF by WB and corresponding densitometric analysis (n=3 per treatment group). Snai1-KD inhibits hypoxia+TGFβ1-induced FMT by reducing Snai1 and LOXL2 expression and colocalization. HCAEC: Human coronary artery endothelial cell. HCF: Human Cardiac Fibroblast.

Snai1-KD inhibited FB-to-MFB transition in HCFs *in vitro* (Fig. 7I, S6) resulted in significant reduction in Snai1 expression (p=0.0002; Fig. 7J) as validated by qPCR. Snai1-KD resulted in a significant reduction of LOXL2 (p=0.0294; Fig. 7K), α-SMA (p=0.0186; Fig. 7L) but increased Vimentin (p=0.0230; Fig. 7M) in HCFs under Hypoxia+TGFβ1 *in vitro*.

Snai1-KD inhibited FB-to-MFB transition as documented by reduced α-SMA in HCFs treated with hypoxia+TGFβ1 by decreasing Snai1 and LOXL2 co-localization in the nuclear and perinuclear areas (Fig. 7N). Immunoblots and corresponding densitometric analysis also showed significantly reduced Snai1 (p=0.0006; Fig. 7O), LOXL2 (p=0.0110; Fig. 7P), α-SMA (p=0.0263; Fig. 7Q) but increased Vimentin (p=0.0071; Fig. 7R) in the Snai1-KD HCFs and further supported our immunofluorescence findings.

## Discussion

In this study, we demonstrate for the first time the molecular mechanisms of EndMT and fibrosis in preclinical and clinical RV failure secondary to pulmonary hypertension. Specifically, we highlight the significant concordance between rat (MCT, SuHx, and PAB) and human RV transcriptome in PH-RVF demonstrating the presence of EndMT as the top common upregulated pathway and Snai1 as the top common upregulated key EndMT gatekeeper transcription factor. Furthermore, we discover the TGFβ1-Snai1-LOXL2 axis and related network as the key regulatory hub underlying RV EndMT and fibrosis. Using state-of-the-art *in vitro* models we demonstrate cell-type specific involvement of Snai1 in mediating EndMT and FMT *via* a LOXL2-dependent mechanism. Finally, we unravel a novel and promising therapeutic role of targeting Snai1 for the rescue of PH-RVF by inhibiting RV EndMT and fibrosis *via* a LOXL2-mediated mechanism (Fig. 8).

**Fig. 8.**
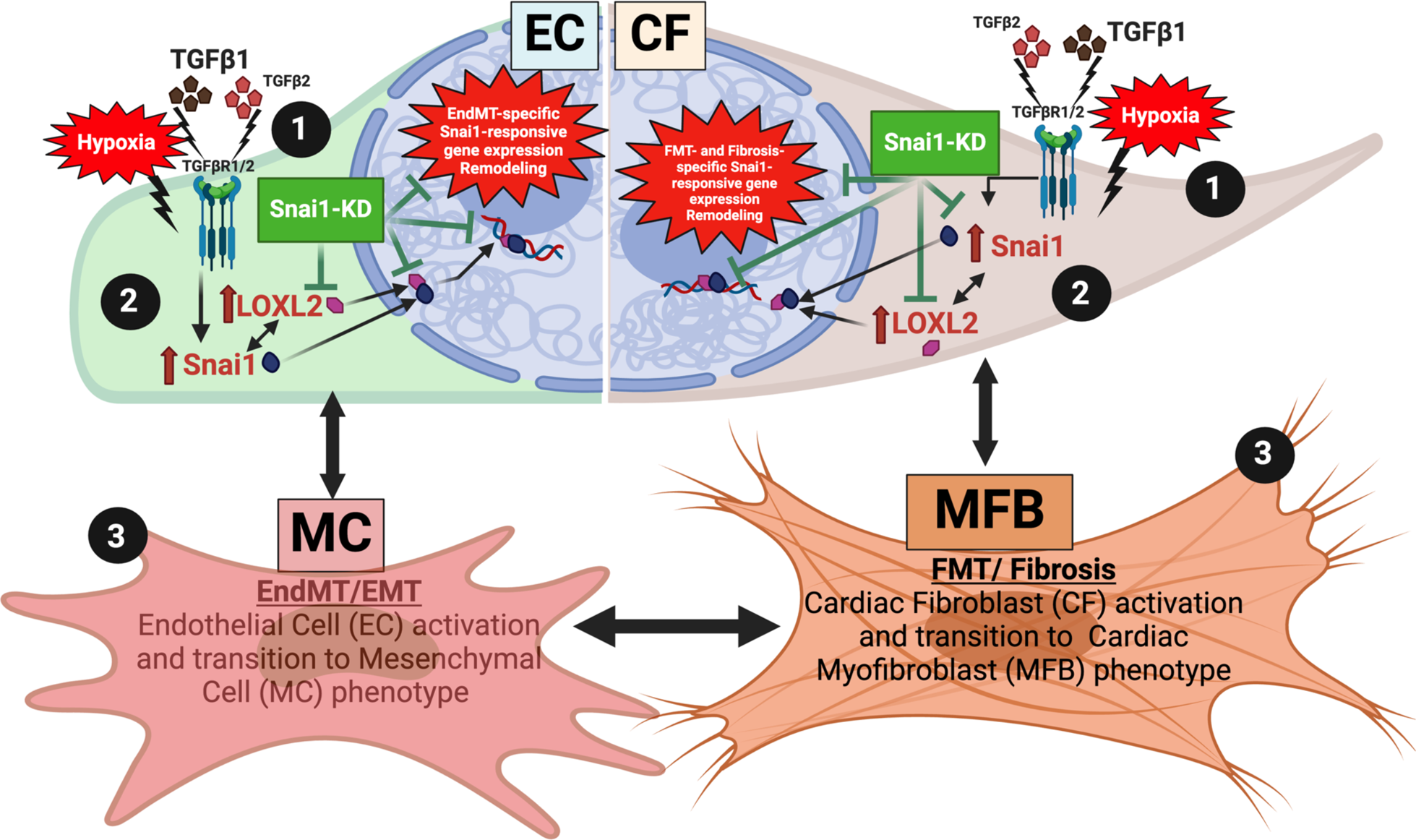
Schematic showing the involvement of TGFβ1-Snai1-LOXL2 axis in inducing RV EndMT, FMT, and fibrosis in PH-RVF and Snai1-KD mediated inhibition of EndMT, FMT, and fibrosis *via* a LOXL2-dependent mechanism. We propose that synergistic stimulation of endothelial cells (EC) and cardiac fibroblasts (CF) by hypoxia+TGFβ1 (depicted by black circle 1) leads to increased Sna1+LOXL2 activation (depicted by black circle 2) resulting in increased EndMT and FMT respectively (depicted by black circle 3). Abbreviations: MC: mesenchymal cell, MFB: myofibroblast, KD: knockdown.

### Role of EndMT and Fibrosis in PH-RVF

Epithelial-to-mesenchymal transition (EMT) *and* endothelial-to-mesenchymal transition (EndMT) play an integral role in early cardiovascular development and are reactivated in numerous chronic cardiovascular disease states ^29,30^. In EndMT, endothelial cells (ECs) acquire a mesenchymal phenotype through a series of molecular events, a gene expression profile similar to smooth muscle cells (SMCs) and develop the ability to migrate and remodel the extracellular matrix (ECM) that promotes their differentiation into SMC-like cells^8^. The hallmarks of EndMT include reduced endothelial markers and increased mesenchymal markers^19^. Recent studies have implicated EMT and EndMT to play an important role in adult cardiovascular diseases and fibrosis. In response to cardiac injury, a significant portion of MFBs arise from the conversion of epithelial or endothelial cells through the EMT or EndMT process, respectively, and ultimately lead to excessive collagen deposition, fibrosis, and cardiac dysfunction^29,31^. EMT and EndMT have been well characterized in myocardial fibrosis and diastolic dysfunction in pressure-overload rodent models^5^ and play a critical role in tissue fibrosis and pulmonary vascular remodeling in PAH^32,33^. However, the precise molecular underpinnings of RV EndMT and fibrosis in PH-RVF remain elusive.

RVF is a significant prognostic determinant of morbidity and mortality in PH^3–7,9,23^. Recent evidence suggests RV fibrosis plays a crucial role in the development of RV failure^23,34,35^. RV fibrosis is associated with increased ventricular stiffness, impaired coronary blood flow, diastolic dysfunction, and arrhythmias^4^. Increased fibrotic deposition, characterized by MFB accumulation and excessive collagen secretion, can result in RV stiffness and severe dysfunction^34^. Based on our Hallmark enrichment analysis of RV from severe rat models (MCT, SuHx, and PAB) of PH-RVF and human decompensated PH-RVF^9,23,24^, we identified up-regulated genes that were highly enriched in functions associated with EMT/EndMT and fibrosis (Fig. 2).

### Role of Snai1 and LOXL2 in EndMT and Fibrosis

Transcription factors such as Snai1/2, TWIST1, and ZEB1/2 are the key EMT and EndMT gatekeepers that interact to culminate in chromatin reorganization and bind to cell-specific gene promoters to cause EMT and EndMT^19^. During EMT and EndMT, Snai1 recruits LOXL2, a well-known transcriptional corepressor, to pericentromeric regions to enable the acquisition of mesenchymal phenotype^20,21^. Snai1 is reported to be present unambiguously in all cardiac cell types^36^. Snai1 and its related network (TGFβ1-Snai1-LOXL2) play a crucial role in promoting EMT/EndMT and fibrosis in various organs^10,28,37–39^. In fact, hypoxia and TGFβ1/2 are known to upregulate Snai1^10,26,27^, which recruits LOXL2 to induce EMT/EndMT^28^. EndMT is known to contribute to cardiac fibrosis by various mechanisms^5^ but its precise role in the progression of RV fibrosis in PH-RVF is yet to be elucidated. Snai1 is critical in mediating trans-differentiation of FBs to MFBs and reprogramming their fibrotic response to further contribute towards ECM deposition and fibrosis progression^37,40–45^. Snai1 mediates cardiac FB-to-MFB transition following TGFβ stimulation^37,46^. Hypoxic injury-induced activation of Snai1 stimulates the expression of fibrosis-related genes and promotes cardiac FBs to attain the MFB fate^37,46^. Snai1-induced activation of MFBs results in deposition of ECM remodeling proteins that can maintain elevated Snai1 expression in the MFBs and help propagate fibrosis^37,40–45^. Although Snai1 and LOXL2 are involved in fibrotic processes^10,39^, however, their precise role in RV EndMT and fibrosis is still not known.

Our comprehensive comparative RNA-Seq analysis shows that EndMT is the common top-most upregulated pathway in the failing RV of rat MCT, SuHx, and PAB models as well as in decompensated human PH-RVF (Fig. 2). Interestingly, from the common EndMT-inducing transcription factors (Snai1, Snai2, Twist1, Zeb1), only Snai1 was preferentially upregulated in RV of MCT, SuHx, and PAB rat models and human decompensated RVF (Fig. 2). The increased expression of Snai1 was associated with TGFβ1, LOXL2 upregulation (Fig. 2).

To highlight the functional importance of Snai1, we performed silencing of Snai1 *in vivo* and *in vitro*. Our *in vivo* data showed that Snai1-KD in the RV not only reversed RV EndMT and fibrosis but also restored RV function and rescued PH in the MCT rats (Fig. 6). More importantly, *in vivo* Snai1-KD also resulted in LOXL2 downregulation in the RV (Fig. 6). Furthermore, our *in vitro* data demonstrated that Snai1-KD resulted in inhibition of EndMT and FB-to-MFB transition *via* a LOXL2-mediated mechanism (Fig. 7).

### Importance of RV Function in PH-RVF and its Link to RV Fibrosis

Growing evidence from preclinical and clinical studies shows that RV fibrosis is detrimental to RV function^15–18^. Interestingly a prior study by Crnkovic^47^ is at odds with the extensive evidence showing that RV fibrosis is deleterious for RV function^15–18,48–52^. This is likely due to the off-target effects of Pirfenidone as discussed by Bogaard et al.^18^ In addition, although a mouse PAB model showed less RV EndMT^53^, our current and published data from multiple rat models of decompensated PH-RVF and human decompensated PAH-RVF with much higher RV pressure overload have consistently shown higher EndMT and fibrosis^9,23,24^. Interestingly, in our study, targeting the TGFβ1-Snai1-LOXL2 axis and manipulating Snai1 resulted in the restoration of RV function by ameliorating EndMT and fibrosis progression in pre-clinical PH-RVF.

### Limitations

Although only male rats were used in this study, future studies on females investigating the mechanistic role of the TGFβ1-Snai1-LOXL2 axis and evaluating the therapeutic potential of Snai1 inhibition are warranted.

## Conclusions

In conclusion, our study highlights significant concordance in RV transcriptome across multiple rat models (MCT, SuHx, PAB) and human decompensated PH-RVF. Using state-of-the-art *in vivo* and *in vitro* models and sophisticated bioinformatics analyses, we demonstrate molecular mechanisms of RV EndMT and fibrosis and discover the novel TGFβ1-Snai1-LOXL2 axis in preclinical and clinical PH-RVF. RV-specific targeting of Snai1 rescues PH-RVF by inhibiting EndMT and fibrosis *via* a LOXL2-mediated mechanism. Targeting Snai1 could serve as a novel and promising RV-specific therapeutic target for PH-RVF.

## Supplementary Figure Legends

**Supplemental Fig. S1.**
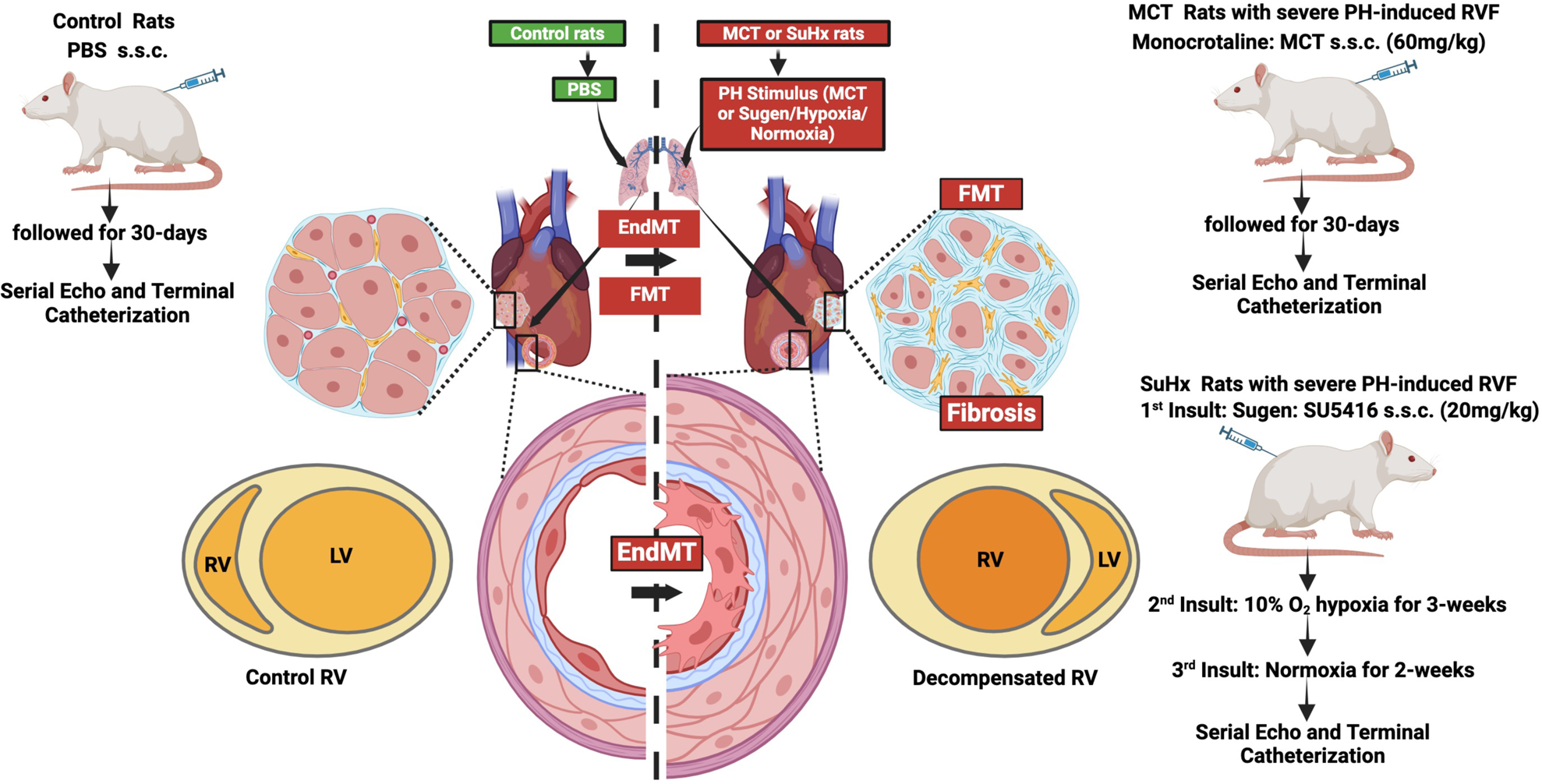
Schematic showing the protocol for induction and development of PH-induced RV failure in MCT and SuHx PH-RVF rats.

**Supplemental Fig. S2.**
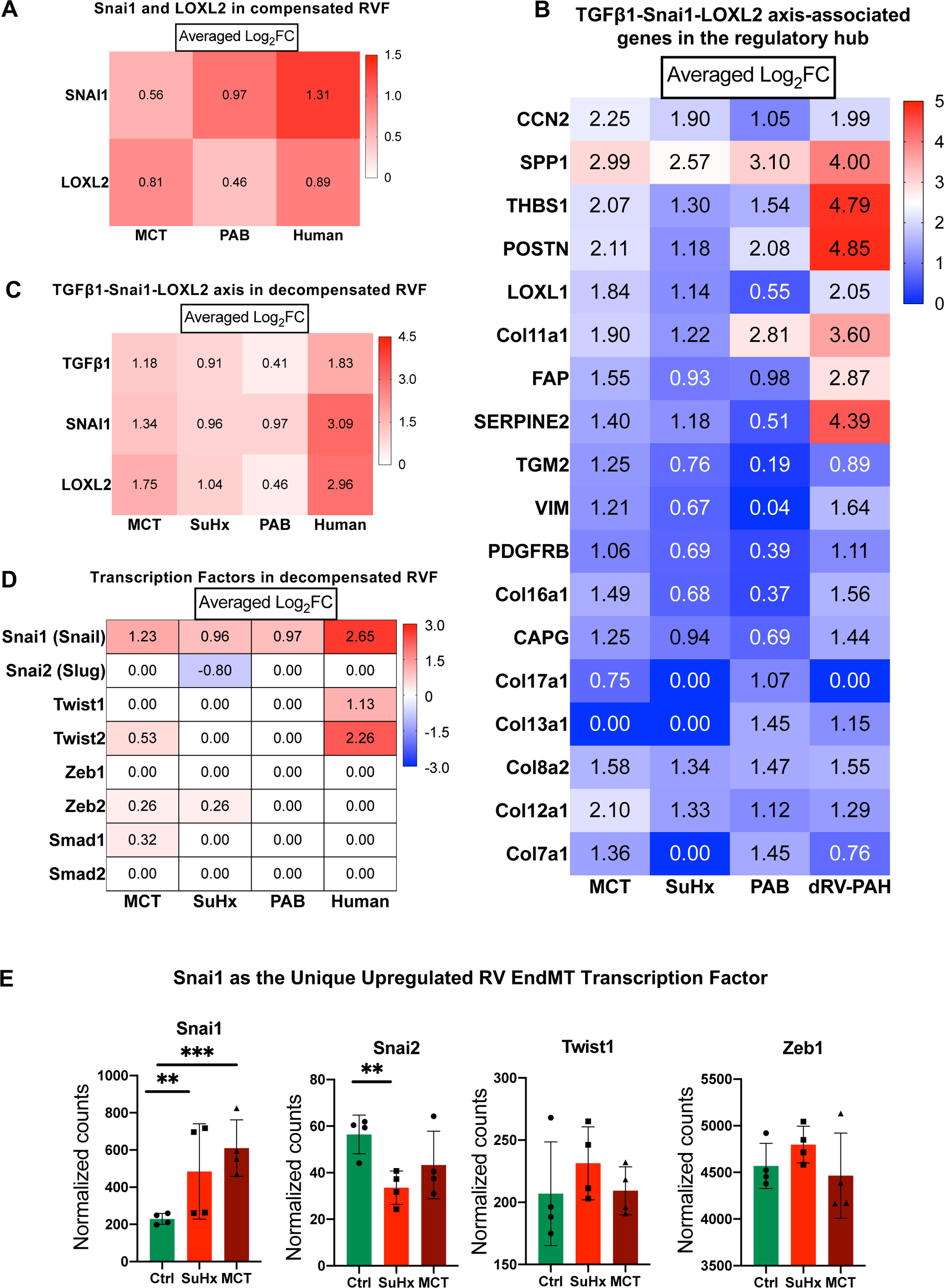
Averaged Log2FC-based Heatmaps of **(A)** Snai1 and LOXL2 in compensated RVs of MCT, PAB rats, and human PH-RVF, Averaged Log2FC-based Heatmaps of **(B)** TGFβ1-Snai1-LOXL2 axis-associated genes in the regulatory hub **(C)** TGFβ1-Snai1-LOXL2 axis, **(D-E)** EndMT-gatekeeper transcription factors in decompensated RVs of MCT, SuHx, PAB rats, and human PH-RVF,

**Supplemental Fig. S3.**
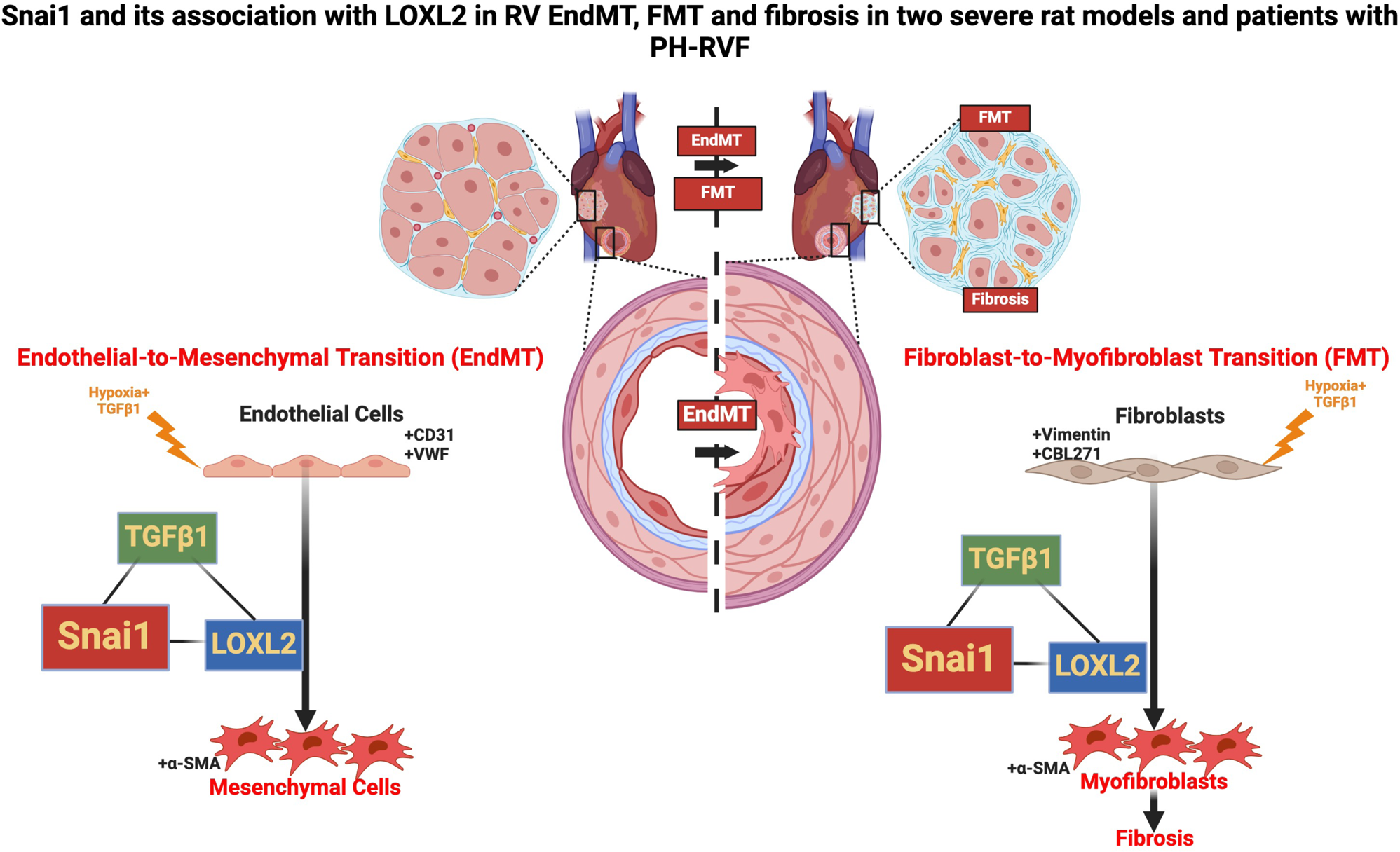
Schematic showing the involvement of TGFβ1-Snai1-LOXL2 axis in inducing RV EndMT, FMT, fibrosis, and development of PH-induced RV failure in MCT and SuHx rats, and human PH-patients.

**Supplemental Fig. S4.**
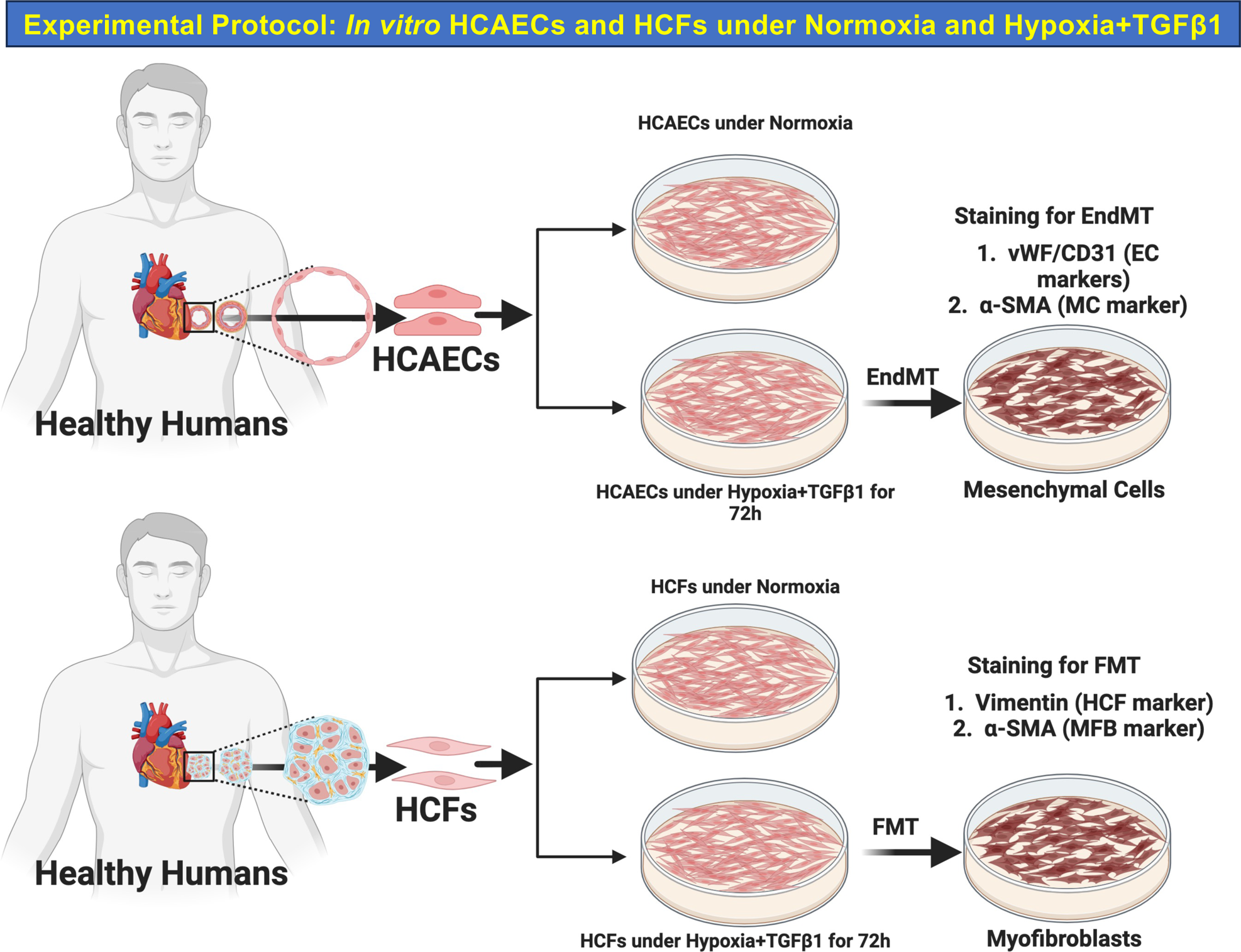
Schematic showing the experimental protocol for inducing EndMT in HCAECs and FMT in HCFs under Hypoxia+TGFβ1 *in vitro*.

**Supplemental Fig. S5.**
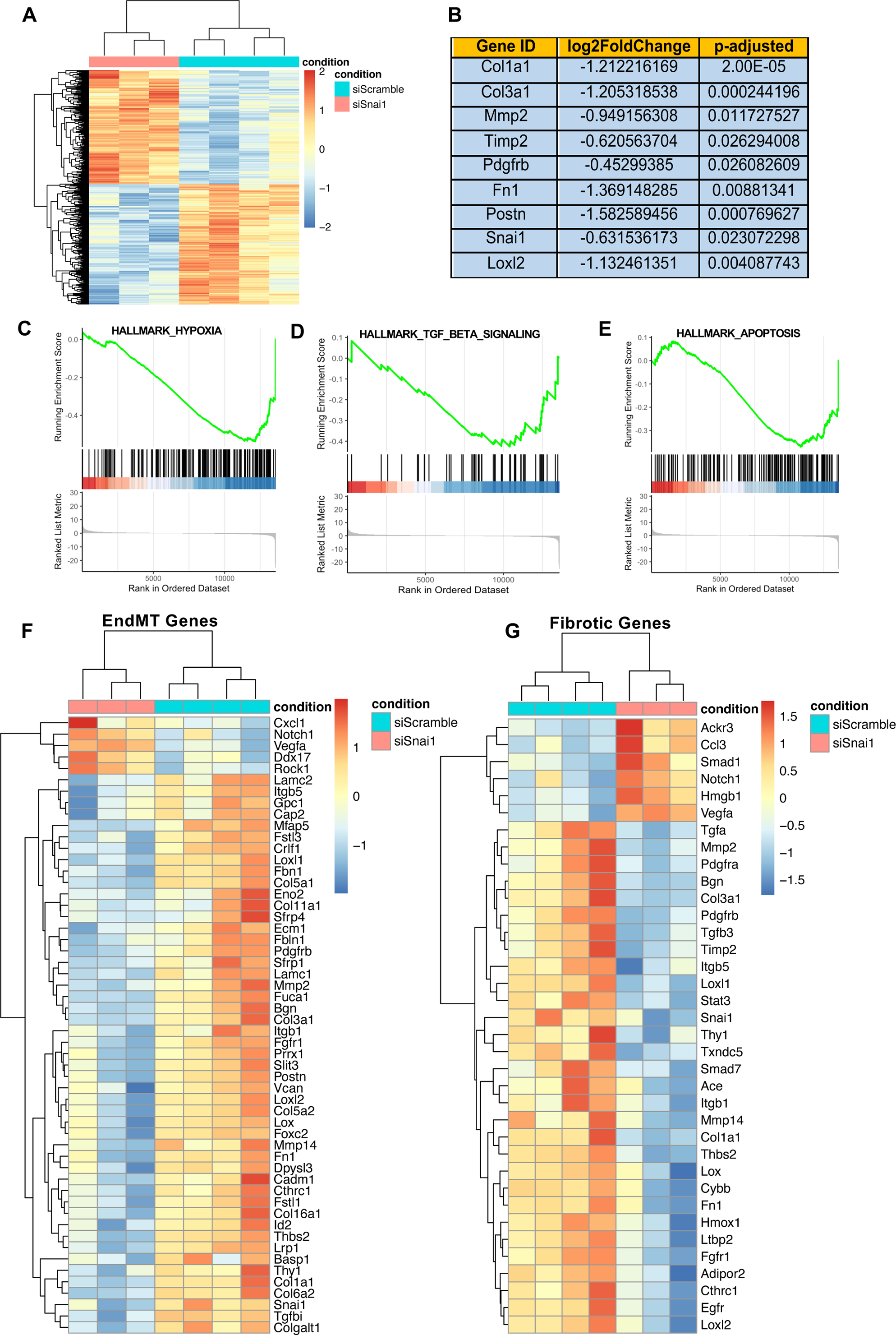
**(A)** Heatmap of the differentially expressed genes, **(B)** Heatmaps of the TGFβ1-Snai1-LOXL2 axis and EndMT-gatekeeper transcription factors, **(C-E)** RV RNASeq Hallmark pathway-specific Gene-set Enrichment analysis showing the downregulation and Snai1-KD mediated successful reversal of hypoxia, TGFβ signaling, and apoptosis, **(F-G)** Hallmark Gene-set Enrichment analysis showing Snai1-KD mediated successful downregulation and reversal of key driver and mediator genes of RV EMT/EndMT, FMT and fibrosis-related pathways in PH-RVF MCT rats. N=3-4 per group. FDR<0.05.

**Supplemental Fig. S6.**
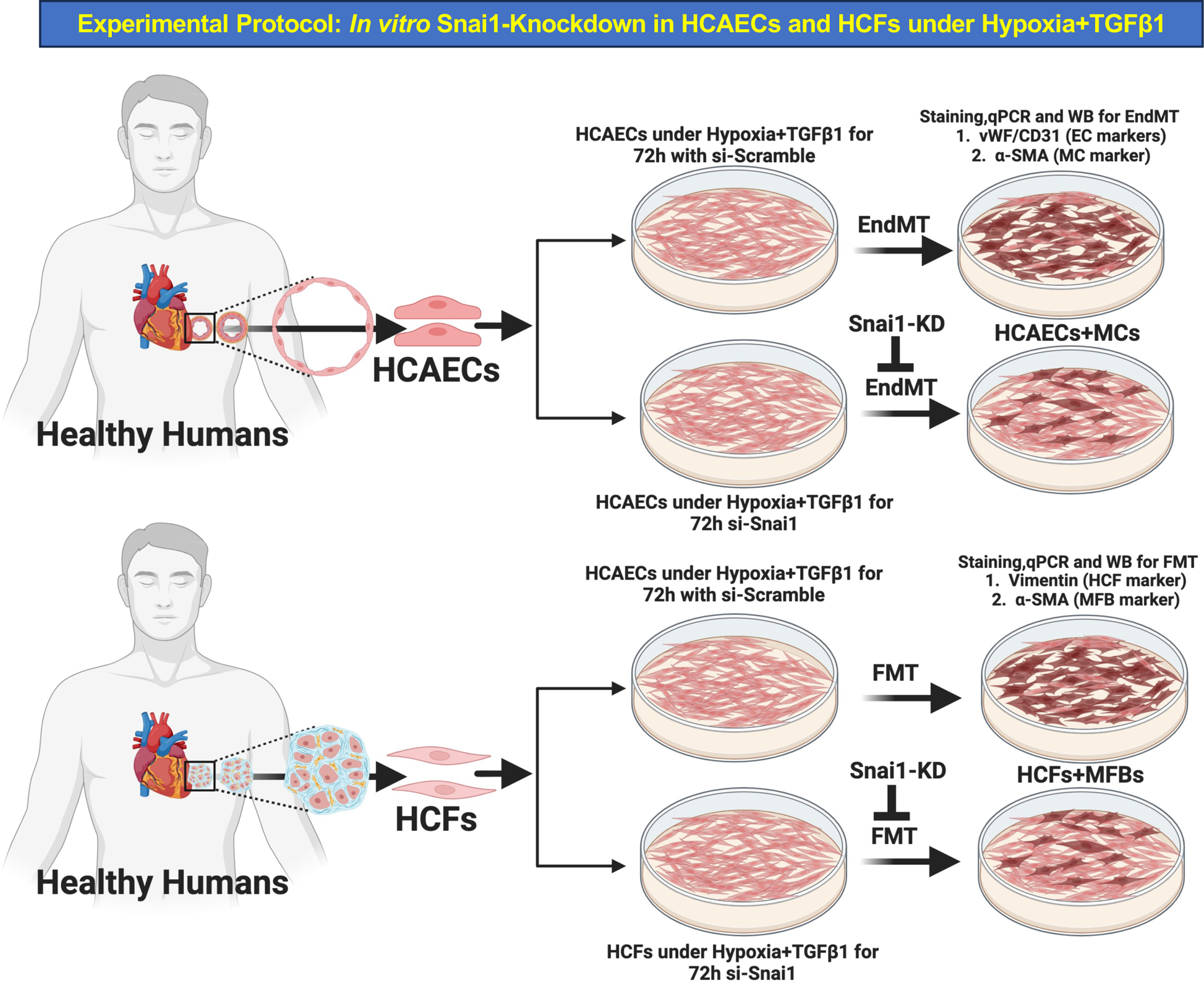
Schematic showing the experimental protocol for Snai1-knockdown in HCAECs and HCFs under Hypoxia+TGFβ1 *in vitro* and assessment of Snai1-KD mediated inhibition of EndMT in HCAECs and FMT in HCFs *via* a LOXL2-dependent mechanism.

---

**Supplemental Table S1:**
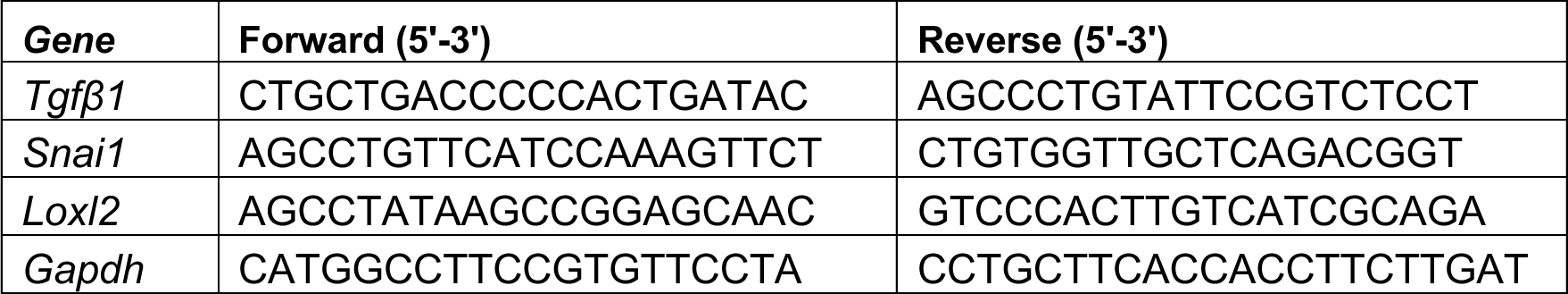
Rat Primers used for RV tissue qPCR.

**Supplemental Table S2:**
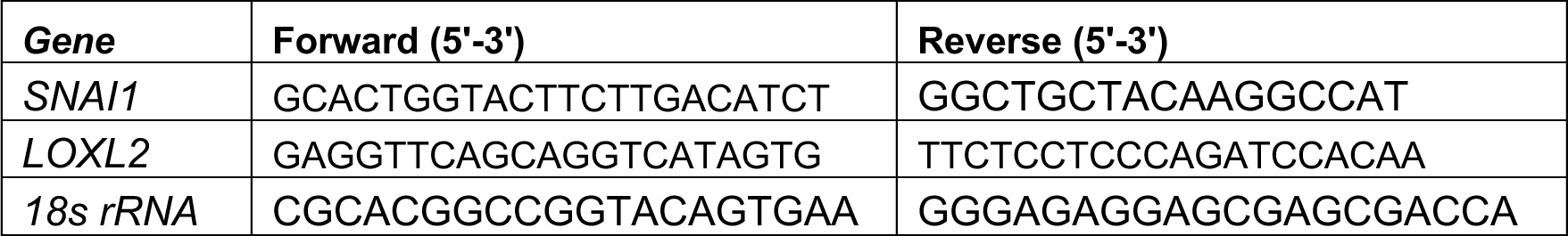
Human Primers used for RV tissue qPCR.

**Supplemental Table S3:** Primary Antibodies used for IF and WB.

| **Antibodies** | **Catalogue No.** | **Vendor** | **IF/WB** | **Dilution** |
| --- | --- | --- | --- | --- |
| Rabbit anti-vWF | 11778-1-AP | Proteintech | IF | 1:200 |
| Rabbit anti-CD31 | NB100-2284 | Novus Biologicals | IF | 1:200 |
| Mouse anti-α-SMA | A2547 | Millipore Sigma | IF | 1:400 |
| Goat anti-Snai1 | PA5-18574 | Invitrogen | IF | 1:200 |
| Rabbit anti-Vimentin | NBP1-31327 | Novus Biologicals | IF | 1:250 |
| Rabbit anti-LOXL2 | NBP1-32954 | Novus Biologicals | IF | 1:200 |
| Mouse anti-LOXL2 | NBP1-32954 | Novus Biologicals | IF | 1:200 |
| Mouse anti-Fibroblasts (CBL271) | CBL271 | Millipore Sigma | IF | 1:200 |
| Goat anti-Vimentin | AF2105 | R&D Systems | IF | 1:200 |
| Mouse anti-α-SMA | A2547 | Millipore Sigma | WB | 1:5000 |
| Rabbit anti-Vimentin | NBP1-31327 | Novus Biologicals | WB | 1:2500 |
| Rabbit anti-Snail (C15D3) Snai1 | 3879 | Cell Signaling Technology | WB | 1:1000 |
| Rabbit anti-LOXL2 | NBP1-32954 | Novus Biologicals | WB | 1:1000 |
| Rabbit anti-GAPDH (14C10) | 2118 | Cell Signaling Technology | WB | 1:5000 |
| Mouse anti-Vinculin | V9131 | Millipore Sigma | WB | 1:5000 |

**Supplemental Table S4:** Secondary Antibodies used for IF and WB.

| **Antibodies** | **Catalogue No.** | **Vendor** | **IF/WB** | **Dilution** |
| --- | --- | --- | --- | --- |
| Alexa Fluor 594 goat anti-mouse IgG (H+L) | A11005 | Invitrogen | IF | 1:1000 |
| Alexa Fluor 594 goat anti-rabbit IgG (H+L) | A11012 | Invitrogen | IF | 1:1000 |
| Alexa Fluor 488 goat anti-mouse IgG (H+L) | A11001 | Invitrogen | IF | 1:1000 |
| Alexa Fluor 488 goat anti-rabbit IgG (H+L) | A11008 | Invitrogen | IF | 1:1000 |
| Alexa Fluor 488 donkey anti-goat IgG (H+L) | A11055 | Invitrogen | IF | 1:1000 |
| Alexa Fluor Cy5 goat anti-mouse IgG (H+L) | A10524 | Invitrogen | IF | 1:1000 |
| Alexa Fluor Cy5 goat anti-rabbit IgG (H+L) | A10523 | Invitrogen | IF | 1:1000 |
| Goat anti-mouse IRDYE 680RD | 926-68070 | LICOR | WB | 1:10000 |
| Goat anti-rabbit IRDYE 680RD | 926-68071 | LICOR | WB | 1:10000 |
| Goat anti-mouse IRDYE 800CW | 926-32210 | LICOR | WB | 1:10000 |
| Goat anti-rabbit IRDYE 800CW | 926-32211 | LICOR | WB | 1:10000 |

**Supplemental Table S5:**
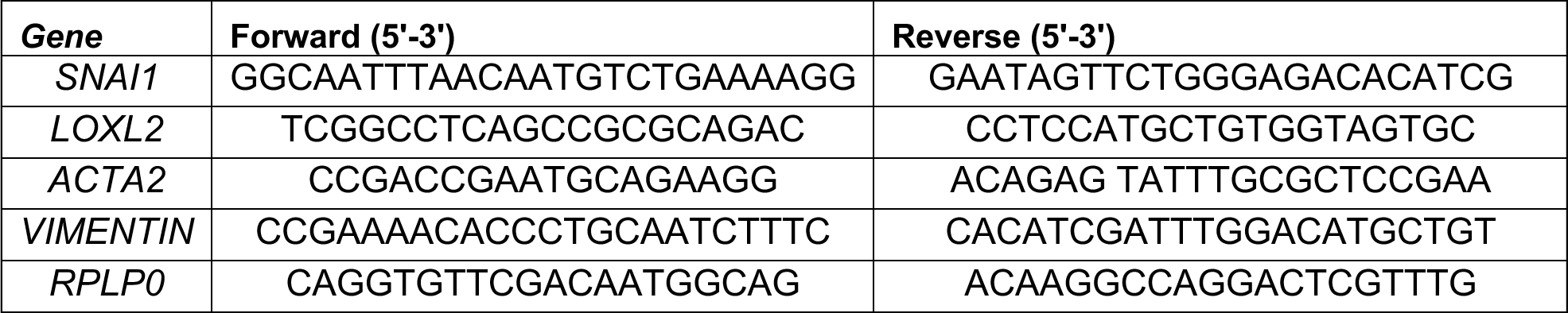
Human Primers used for cell culture qPCR.

**Supplemental Table S6:** Patients and control subjects demographics.

| **Human Sample** | **Age** | **Sex** |
| --- | --- | --- |
| Control 1 | 28 | M |
| Control 2 | 56 | M |
| Control 3 | 53 | M |
| Control 4 | 29 | M |
| Control 5 | 47 | M |
| Control 6 | 59 | M |
| Control 7 | 50 | M |
| Control 8 | 22 | M |
| Control 9 | 58 | M |
| Control 10 | 46 | M |
| PAH 1 (SSc-PAH) | 77 | M |
| PAH 2 (SSc-PAH) | 47 | F |
| PAH3 (IPAH) | 74 | F |
| PAH 4 (LV-HTP) | 57 | F |
| PAH 5 (Portal-PAH) | 46 | F |
| PAH 6 (Familial-PAH) | 61 | F |
| PAH 7 (IPAH) | 65 | M |
| PAH 8 (VOD-PAH) | 72 | F |
| PAH 9 (SSc-PAH) | 62 | M |
| PAH 10 (SSc-PAH) | 53 | F |

